# Lipid Nanoparticle-Associated Inflammation is Triggered by Sensing of Endosomal Damage: Engineering Endosomal Escape Without Side Effects

**DOI:** 10.1101/2024.04.16.589801

**Authors:** Serena Omo-Lamai, Yufei Wang, Manthan N. Patel, Eno-Obong Essien, Mengwen Shen, Aparajeeta Majumdar, Carolann Espy, Jichuan Wu, Breana Channer, Michael Tobin, Shruthi Murali, Tyler E. Papp, Rhea Maheshwari, Liuqian Wang, Liam S. Chase, Marco E. Zamora, Mariah L. Arral, Oscar A. Marcos-Contreras, Jacob W. Myerson, Christopher A. Hunter, Andrew Tsourkas, Vladimir Muzykantov, Igor Brodsky, Sunny Shin, Kathryn A. Whitehead, Peter Gaskill, Dennis Discher, Hamideh Parhiz, Jacob S. Brenner

## Abstract

Lipid nanoparticles (LNPs) have emerged as the dominant platform for RNA delivery, based on their success in the COVID-19 vaccines and late-stage clinical studies in other indications. However, we and others have shown that LNPs induce severe inflammation, and massively aggravate pre-existing inflammation. Here, using structure-function screening of lipids and analyses of signaling pathways, we elucidate the mechanisms of LNP-associated inflammation and demonstrate solutions. We show that LNPs’ hallmark feature, endosomal escape, which is necessary for RNA expression, also directly triggers inflammation by causing endosomal membrane damage. Large, irreparable, endosomal holes are recognized by cytosolic proteins called galectins, which bind to sugars on the inner endosomal membrane and then regulate downstream inflammation. We find that inhibition of galectins abrogates LNP-associated inflammation, both *in vitro* and *in vivo*. We show that rapidly biodegradable ionizable lipids can preferentially create endosomal holes that are smaller in size and reparable by the endosomal sorting complex required for transport (ESCRT) pathway. Ionizable lipids producing such ESCRT-recruiting endosomal holes can produce high expression from cargo mRNA with minimal inflammation. Finally, we show that both routes to non-inflammatory LNPs, either galectin inhibition or ESCRT-recruiting ionizable lipids, are compatible with therapeutic mRNAs that ameliorate inflammation in disease models. LNPs without galectin inhibition or biodegradable ionizable lipids lead to severe exacerbation of inflammation in these models. In summary, endosomal escape induces endosomal membrane damage that can lead to inflammation. However, the inflammation can be controlled by inhibiting galectins (large hole detectors) or by using biodegradable lipids, which create smaller holes that are reparable by the ESCRT pathway. These strategies should lead to generally safer LNPs that can be used to treat inflammatory diseases.

## Introduction

Lipid nanoparticles (LNPs) have emerged as the biopharmaceutical industry’s dominant delivery system for RNA delivery, following their successful use in > 1 billion people with the COVID-19 vaccine^1^. Subsequently, industry has invested tens of billions of dollars in moving LNPs into late stages of development not just for vaccines, but also for delivering RNA therapeutics for diseases ranging from common cancers to rare monogenic diseases^2,3^. LNPs achieve efficient delivery of RNA, which itself is an incredibly versatile class of therapeutic cargo, allowing for down- or up-regulation of proteins (via siRNA and mRNA, respectively) and for gene-editing. In addition to efficient delivery of RNA, the other reason for the widespread use of LNPs is their safety record, which is well illustrated by the COVID-19 vaccines^4,5^.

However, LNPs’ clear safety in the COVID-19 vaccines does not necessarily extend to situations in which LNPs are delivered via different routes and for non-vaccine indications. Most importantly, our group and others have recently shown that LNPs can induce severe inflammation and worsen markers of pre-existing inflammation by up to>10-fold^6–11^. LNPs have been shown to activate the innate immune system across multiple routes of administration, including intranasally, intramuscularly, and intravenously. They induce the secretion of pro-inflammatory cytokines, such as IL-6 and TNF-⍺, and chemokines that promote infiltration of activated leukocytes that damage tissues. If LNPs’ pro-inflammatory effects are not understood mechanistically and mitigated by re-engineering, LNPs will likely fail in patients with pre-existing inflammatory diseases (including most elderly and hospitalized patients), and certainly will be unable to safely treat common inflammatory diseases, such as heart attack and stroke. In fact, before treatment with patisiran, (which was approved by the FDA in 2018 as the first LNP nucleic acid-loaded therapeutic), patients were dosed with immunosuppressants such as the anti-inflammatory steroid dexamethasone to prevent side effects^12^. Thus, LNPs’ current highly inflammatory nature significantly worsens their safety profile and is a major roadblock to LNP therapeutics^10^

In retrospect, LNP-associated inflammation could have been anticipated, as LNPs are one of the best-known adjuvants for vaccines, and adjuvanticity shares many mechanisms with inflammation. Even empty LNPs (no RNA cargo) induce comparable B- and T-cell responses to traditional vaccine adjuvants^13–16^. Empty LNPs co-delivered with recombinant SARS-CoV2 spike protein provide vaccine responses roughly equivalent to those of LNPs containing mRNA encoding the spike protein, showing that the COVID-19 vaccine owes much of its power to the LNPs’ adjuvant effect^17^. Thus, to further improve vaccines and add to their immunogenicity while minimizing their reactogenicity (unwanted local and remote inflammation), we contend that we must understand how LNPs achieve their adjuvanticity^18^.

Therefore, we sought to understand the fundamental mechanisms by which LNPs induce inflammation, exacerbate pre-existing inflammation, and provide adjuvanticity, all of which are critical to the future of LNP and RNA medicines. We began with the initial hypothesis that LNPs’ greatest strength, endosomal escape, is a double-edged sword and directly activates inflammation. Endosomal escape occurs when ionizable lipids in LNPs interact with the endosomal membrane, creating a pore or otherwise translocating the LNPs’ cargo mRNA from the endosomal lumen into the cytosol. Endosomal escape occurs when ionizable lipids in LNPs interact with the endosomal membrane, creating a pore or other means of translocating the LNPs’ cargo RNA from the endosomal lumen into the cytosol. Our hypothesis therefore posits that the cell detects leakage of endosomal content, escaping alongside cargo RNA into the cytosol, and this activates inflammation.

To investigate this hypothesis, we took two complementary approaches. First, we performed structure-function studies to find LNP features that affect inflammation. Second, we investigated upstream signaling pathways leading from LNPs to inflammation. We found that, as hypothesized, LNPs’ endosomal escape induces endosomal membrane damage directly activates inflammation. However, the degree of inflammation is not just a function of the *quantity* of endosomal holes, but also the *size* of endosomal holes. The primary sensors of large endosomal ruptures are sugar-binding lectins, known as galectins, which detect the exposure of glycans to the cytosol upon membrane damage^19,20^. When LNP-induced endosomal ruptures are smaller, cells recruit the ESCRT machinery to promote endosomal repair^21–24^. We show that LNPs with a particular ionizable lipid create intermediate-sized holes, marked by ESCRT. These LNPs drive strong expression from cargo mRNA while not inducing inflammation. Further, we show that inhibiting galectins, and therefore large endosomal damage sensing, ameliorates LNP-induced inflammation *in vitro* and *in vivo* across multiple routes of delivery. Finally, we show LNPs designed to create ESCRT-recruiting endosomal holes can avoid LNP-induced exacerbation of inflammation and can ameliorate a highly inflammatory disease model with delivery of therapeutic mRNA. These findings show that LNP-induced membrane damage produces both RNA endosomal escape and inflammation, but such endosomal damage can be controlled to provide high-expressing, non-inflammatory LNPs.

## Results

### LNPs induce potent inflammation *in vitro* and *in vivo* across species

We first probed the inflammation induced in various systems by mRNA-LNPs fabricated with one of the best-studied ionizable lipids, cKK-E12, which is among the top-reported ionizable lipids for driving strong RNA expression^25^ (Supplementary Table 1). Into healthy, wildtype mice, we intratracheally instilled LNPs containing an mRNA dose ranging from 2.5µg - 10µg per mouse, which are therapeutically relevant doses for intratracheal LNP treatment^26,27^. Animals were sacrificed 24h after LNP treatment. Gross anatomical inspection of the lungs from mice instilled with 7.5µg of mRNA in LNPs reveals severe “hepatization,” which is the classical pathology term for lung inflammation that makes lung tissue resemble liver tissue (Fig. 1A). We assessed lung-specific inflammation by analyzing protein and leukocyte content in the bronchoalveolar lavage (BAL) fluid, which assess capillary leak and leukocyte infiltration into the alveoli (air sacs). We observed large, dose-dependent increases in BAL protein (Fig. 1B) and BAL leukocyte (Fig. 1C) levels from 2.5µg - 10µg of mRNA. We also observed dose-dependent changes in other markers associated with inflammation (Supplementary Fig. 1). To put this inflammation severity into context, we gave mice nebulized LPS, which is the most common animal model of acute respiratory distress syndrome (ARDS), the acute alveolar inflammation caused by severe COVID-19, sepsis, and other insults. Mice given nebulized LPS had lower BAL inflammatory markers than intermediate doses of mRNA-LNPs (Supplementary Fig. 2A).

**Figure 1:**
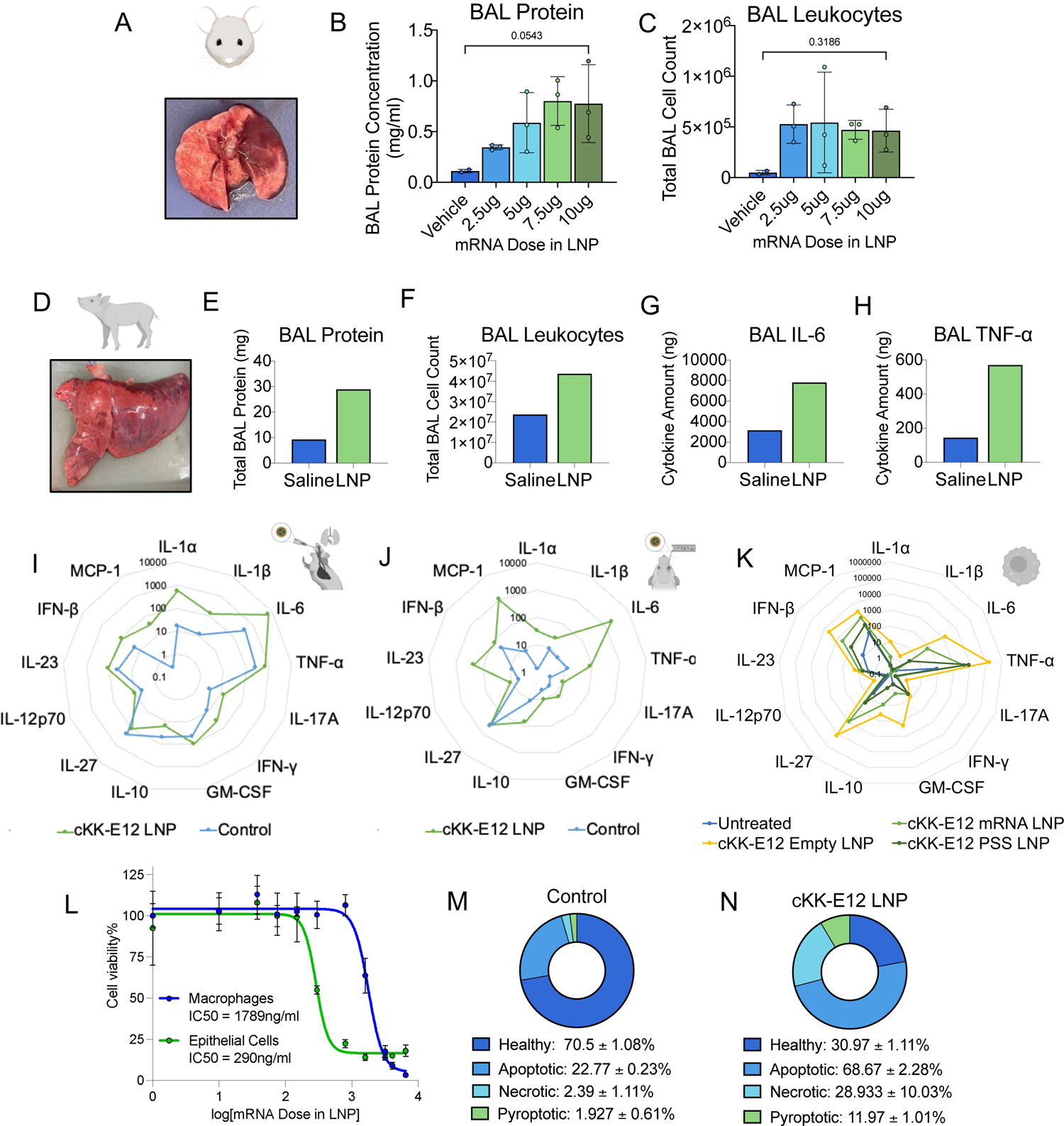
LNPs induce inflammation *in vitro* in various cell types and *in vivo* across species when administered intratracheally (IT) or intravenously (IV). (A) Gross anatomical image of lungs from mice intratracheally instilled with 7.5µg of mRNA in cKK-E12 LNPs per mouse. LNPs were instilled intratracheally into mice and lungs were harvested 24h after. Effect of the dose of intratracheally instilled cKK-E12 LNPs on (B) protein levels and (C) leukocyte count in the bronchoalveolar lavage (BAL) fluid 24 post-LNP instillation. LNPs increase BAL protein and leukocytes in a dose-dependent manner. (D) Gross anatomical image of *ex vivo* pig lungs instilled with cKK-E12 LNPs. LNPs or saline were incubated in the lungs for 3h at 37℃ at a dose of 0.8mg of mRNA in LNPs. Compared to control *ex vivo* pig lungs instilled with saline, the BAL fluid extracted from LNP-treated lungs shows increased levels of (E) protein, (F) leukocytes, (G) IL-6, and (H) TNF-α. (I) Multiplex analysis of cytokine and chemokine concentrations (IL-1α, IL-1β, IL-6, TNF-α, IL-17A, IFN-γ, GM-CSF, IL-10, IL-27, IL-12p70, IL-23, IFN-β, MCP-1) induced in the BAL fluid by intratracheally instilled cKK-E12 LNPs. LNPs were intratracheally instilled at a dose of 7.5µg and BAL fluid was extracted 2h later. (J) Cytokine and chemokine concentrations in the plasma after intravenous injection of cKK-E12 LNPs. LNPs were injected at a dose of 7.5µg of mRNA in LNPs per mouse and allowed to circulate for 2h. (K) Cytokine and chemokine concentrations induced in RAW 264.7 macrophages by cKK-E12 mRNA LNPs, cKK-E12 empty LNPs (formulated with no cargo), and cKK-E12 LNPs formulated with a negatively charged polymer, polystyrene sulfonate (PSS). Cells were treated with LNPs for 6h and the supernatant was collected for multiplex analysis. (L) Effect of cKK-E12 LNP dose on cell viability after 24h in RAW 264.7 macrophages and MLE-12 epithelial cells. Fraction of cell death induced by apoptosis, necrosis, or pyroptosis in (M) control RAW 264.7 macrophages and (N) RAW 264.7 macrophages treated with 400ng/ml of mRNA in cKK-E12 LNPs 24h after treatment. After LNP treatment, cells were isolated and stained with various markers for apoptosis, necrosis, and pyroptosis for flow cytometry analysis. Statistics: For (E) – (H), n=3 replicates from 1 *ex vivo* pig lung. For all other graphs, n=3 and the data shown represents mean ± SEM. For (A) & (B), comparisons between graphs were made using one-way ANOVA with Tukey’s post-hoc test.

To generalize the inflammation stimulated by LNPs to other species, we instilled cKK-E12 LNPs or saline into *ex vivo* pig lungs for 3 hours at a dose of 0.8mg of mRNA in LNPs and subsequently extracted the BAL fluid. Similarly to mice, we observed hepatization after LNP administration (Fig. 1D). We observed a > 3-fold increase in BAL protein levels (Fig. 1E) and a > 1.8-fold increase in BAL leukocyte count (Fig. 1F) relative to saline-treated control samples. The BAL concentrations of the pro-inflammatory cytokines IL-6 (Fig. 1G) and TNF-⍺ (Fig 1H) increased by ∼2.5-fold and ∼4-fold respectively.

In mice, we investigated more deeply the cytokine profile post-mRNA-LNP treatment *in vivo* and *in vitro*. We intratracheally instilled LNPs in mice at a dose of 7.5µg of mRNA in LNPs, waited two hours, and harvested BAL fluid. We observed a significant increase in the BAL concentrations of pro-inflammatory cytokines (particularly IL-1α, IL-6, TNF-α, IFN-β) and chemokines (MCP-1) (Fig 1I). Although the magnitudes differed, we observed an upregulation of the same cytokines and chemokines in plasma 2 hours after intravenous LNP injection in mice at the same dose (Fig. 1J), showing a common cytokine response profile across different tissues and delivery routes that localize LNPs to different cell types.

We have previously shown that intravenous LNPs worsen pre-existing inflammation in mice, and this effect is abrogated by removal of phagocytes^6^. Therefore, we tested for LNP-induced cytokine profile changes in a macrophage-derived cell line (RAW 264.7) and looked for similarities with the cytokine profile changes induced by LNPs *in vivo*. We exposed RAW 264.7 macrophages to cKK-E12 mRNA-LNPs at a dose of 400 ng/ml for 6h. To isolate the inflammatory effects of the lipid component from that of the mRNA cargo, we also administered the same lipid dose of empty LNPs with no cargo, and LNPs loaded with a negatively charged polymer, polystyrene sulfonate (PSS), as a non-RNA cargo that mimics the negative charge of mRNA^28^. We found that mRNA, PSS, and empty LNPs all upregulate the pro-inflammatory cytokines and chemokines that are upregulated in vivo studies, primarily IL-1α, IL-6, TNF-α, IFN-β and MCP-1 (Fig. 1K). However, empty LNPs lead to the highest cytokine concentrations, inducing a ∼20-fold and a ∼500-fold higher IL-6 concentration than mRNA and PSS LNPs respectively. These results indicate that the inflammation stimulated by LNPs is primarily due to the lipid component, rather than the RNA cargo, as LNP-induced inflammation is actually worsened by the absence of RNA.

To determine how cell types other than macrophages respond to LNPs, we compared the effects of LNP treatment on the viability of RAW 264.7 macrophages vs. an epithelial cell line (MLE-12) (Fig. 1L). Cells were treated with varying doses of cKK-E12 LNPs for 24 hours and the fraction of viable cells was quantified. While there was a significant decrease in cell viability with increasing LNP dose in RAW macrophages, there is a much larger reduction in cell viability in MLE-12 epithelial cells. We measure IC-50 values of 1789 ng/ml for macrophages and 290 ng/ml for epithelial cells. Indeed, epithelial cell viability following LNP treatment was too low to reliably detect an LNP-induced cytokine response (Supplementary Fig. 2B).

We further probed the mechanisms of cell death in macrophages, by analyzing the fraction of cells that were apoptotic, necrotic, or pyroptotic 24h after LNP treatment at a dose of 400ng/ml of mRNA in LNPs. Apoptosis is a type of programmed cell death which is typically non-inflammatory^29^. On the other hand, necrosis and pyroptosis are inflammatory cell death mechanisms that are typically triggered by membrane rupture and inflammasome assembly, respectively^29^. While most cells were healthy under control conditions (Fig. 1M), LNP treatment increased the fraction of apoptotic, necrotic, and pyroptotic cells by 3-fold, 13-fold, and 6-fold respectively (Fig. 1N). Therefore, LNPs have variable effects on different cell types, with LNPs inducing strong cytokine responses and inflammatory forms of cell death in macrophages.

### LNP-induced inflammation depends on the ionizable lipid and positively correlates with mRNA expression

To probe the mechanisms of LNP-induced inflammation, we varied the LNPs’ ionizable lipids. Ionizable lipids facilitate the process of endosomal escape of LNPs’ RNA cargo, enabling transfection ^30^. Once protonated in the acidic endosome, ionizable lipids form ion pairs with the anionic lipids in the endosomal membrane, leading to the formation of an inverse hexagonal phase that disrupts the endosomal membrane and releases the RNA cargo into the cytosol^31–33^. Different ionizable lipids have different endosomal disruption capabilities as a function of lipid structural properties which influence their ability to form the inverse hexagonal phase^34,35^. We hypothesized that these differences also correlate with inflammatory responses elicited by LNPs. We propose that LNPs formulated with less potent ionizable lipids induce less endosomal escape of RNA, lower RNA expression, and less inflammation, and more potent ionizable lipids induce higher RNA expression but more severe inflammatory responses (Fig. 2A). This is part of our central hypothesis that some aspect of endosomal escape itself induces inflammation and follows from our results in the previous section showing that lipid components of LNPs, rather than RNA cargo, drive inflammation (Fig. 1K).

**Figure 2:**
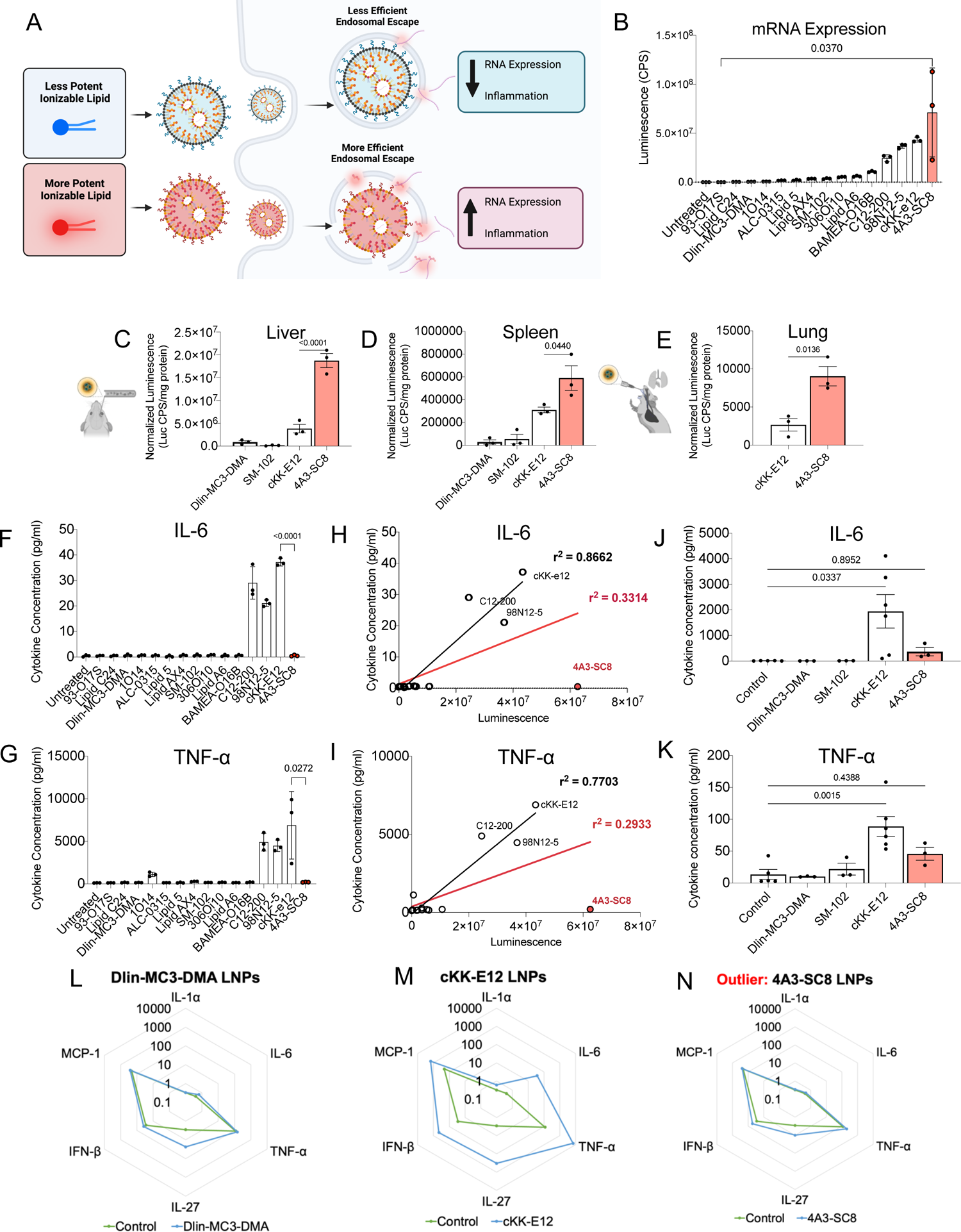
LNP-Induced inflammation is ionizable lipid dependent and positively correlates with mRNA expression. (A) Schematic of hypothesis: LNPs formulated with mild ionizable lipids have less efficient endosomal escape leading to lower mRNA expression and lower inflammation. LNPs formulated with potent ionizable lipids have the reverse effect. (B) Luciferase expression in RAW macrophages of 15 LNPs formulated with 15 different ionizable lipids 6h after treatment. The highest expressing ionizable lipids, cKK-E12 and 4A3-SC8 (red bar) expressed >400-fold and >700-fold more respectively than the lowest expressing ionizable lipid 93-O17S. The LNP luciferase expression profile in vivo in the (C) liver and (D) spleen 6h after intravenous injection into mice follows the in vitro trend Dlin-MC3-DMA LNPs < SM-102 LNPs < cKK-E12 LNPs < 4A3-SC8 LNPs. (E) Similarly, 4A3-SC8 LNPs have a 3-fold higher luciferase expression in the lung than cKK-E12 LNPs 6h after intratracheal administration. In vitro (F) IL-6 and (G) TNF-⍺ concentrations 6h after treatment with the 15 ionizable lipid LNP formulations in (B). In vitro (H) IL-6 and (I) TNF-⍺ concentrations have a positive correlation with luciferase expression. LNPs formulated with the highest expressing ionizable lipids (cKK-E12, C12-200, 98N12-5) are also the most inflammatory except for 4A3-SC8 which does not increase cytokine levels above control levels. This is illustrated in the linear regression fits excluding 4A3-SC8 LNPs (black trendlines, (H) R^2^ = 0.8662 and (I) R^2^ = 0.7703) versus those including 4A3-SC8 LNPs (red trendlines, (H) R^2^ = 0.3314 and (I) R^2^ = 0.2933). After IV LNP injection, the plasma concentrations of (J) IL-6 and (K) TNF-⍺ follow the trend Dlin-MC3-DMA LNPs< SM-102 LNPs < cKK-E12 LNPs in line with the luciferase expression trend. 4A3-SC8 LNPs which do not cause significant cytokine upregulation. Plasma was extracted 2h after LNP treatment. 6h after treatment of *in vitro* RAW macrophages with LNPs, (L) Dlin-MC3-DMA LNPs, which have low mRNA expression, do not increase pro-inflammatory cytokine (IL-1α, IL-6, TNF-α, IL-27, IFN-β, MCP-1) concentrations above control levels, (M) cKK-E12 LNPs, which have high mRNA expression, significantly upregulate pro-inflammatory cytokines, and (N) high-expressing 4A3-SC8 LNPs do not upregulate cytokine levels above control. Statistics: n=3-6 and the data shown represents mean ± SEM. For (H) & (I), analyses were made using a simple linear regression model. For all other graphs, comparisons between groups were made using one-way ANOVA with Tukey’s post-hoc test.

We screened LNPs formulated with varying ionizable lipids and compared their ability to translate cargo RNA (a measure of endosomal escape) to their induction of inflammation. Our primary screen evaluated of 15 of the most studied and potent ionizable lipids^25,36^. Our chosen lipids represent 4 key classes: unsaturated (e.g. Dlin-MC3-DMA), multi-tail (e.g C12-200), biodegradable (e.g. SM-102) and branched tail (e.g 98N12-5^37^) (Supplementary Table 1)^38^. We first confirmed that the uptake of LNPs in RAW macrophages was similar regardless of ionizable lipid (Supplementary Fig. 3). Comparing mRNA expression capacities, we found that luciferase expression induced by these LNPs varied widely, with a ∼700-fold difference between the lowest expressing (93-O17S, Lipid C24, Dlin-MC3-DMA) and the highest expressing (cKK-E12, 4A3-SC8) ionizable lipids (Fig 2B). 4A3-SC8 induced the highest luciferase expression in RAW macrophages^39,40^. We tested a subset of our LNPs intravenously and intratracheally in mice. Matching the expression profile *in vitro*, luciferase expression in the liver and spleen after intravenous injection follows the trend Dlin-MC3-DMA LNPs < SM-102 LNPs << cKK-E12 LNPs < 4A3-SC8 LNPs (Fig 2C, D). 4A3-SC8 LNPs induce >3-fold higher lung luciferase expression than cKK-E12 LNPs after intratracheal administration (Fig. 2E).

We assessed provocation of inflammation by the same library of LNPs, with the goal of correlating luciferase expression and inflammatory profiles. We measured the effect of our LNPs on cell viability in RAW macrophages (Supplementary Fig. 4) and A549 cells (Supplementary Fig. 5). We measured the concentrations of the cytokines IL-6 (Fig. 2F), TNF⍺ (Fig. 2G), IL-1α (Supplementary Fig. 6A), and MCP-1 (Supplementary Fig. 6B) produced by RAW macrophages 6h after treatment with each of the LNPs in our library at a dose of 400ng/ml of mRNA in LNPs. We found that the highest expressing ionizable lipids (C12-200, 98N12-5, cKK-E12) tend to generate the largest cytokine response. To obtain a more detailed structure-function relationship, we expanded on our library by producing a series of lipidoids with the same amine head group with one-carbon differences in their tail structures^41^. For LNPs formulated with these lipidoids, we again found a positive correlation between expression and TNF⍺ upregulation in RAW macrophages (Supplementary Fig. 7). The highest expressing ionizable lipid in our primary screen, 4A3-SC8, breaks this trend, producing cytokine levels matching untreated controls. In scatter plots of cytokine vs. luciferase expression, there is a positive correlation between luciferase expression and inflammatory cytokines if we omit data for 4A3-SC8 LNPs (black trendlines, with r^2^ = 0.8662, 0.7703, 0.7711, and 0.819 for Fig. 2H, Fig. 2I, Supplementary Fig. 6C, and Supplementary Fig. 6D, respectively). The correlation is broken by the inclusion of 4A3-SC8 LNPs (red trendlines, r2 = 0.3314, 0.2933, 0.2726, and 0.3318 for Fig. 2H, Fig. 2I, Supplementary Fig. 6C, and Supplementary Fig. 6D, respectively).

We find similar trends in vivo with intravenous injection of LNPs in mice: 2 hours post-LNP injection, the plasma concentrations of IL-6 (Fig. 2J), TNF⍺ (Fig. 2K), IL-1α (Supplementary Fig. 6E), and MCP-1 (Supplementary Fig. 6F) follow the trend Dlin-MC3-DMA LNPs < SM-102 LNPs < cKK-E12, matching the trends for luciferase expression in the liver and spleen (Fig. 2C-D). As with *in vitro* experiments, 4A3-SC8 LNPs break the trend, causing no significant increase in plasma cytokine levels despite inducing the highest liver and spleen luciferase expression. We also compared cKK-E12 and 4A3-SC8 LNPs in human macrophages and found that cKK-E12 LNPs lead to significantly higher translocation of NF-κB from the cytoplasm to the nucleus (Supplementary Fig. 8 & Supplementary Table 2).

For all but one member of our lipid library, the hypothesis of Figure 2A is correct, in that more strongly expressing ionizable lipids also drive greater inflammation. However, 4A3-SC8 breaks this trend by inducing high expression with no inflammation. In the following section, we elucidate mechanisms by which 4A3-SC8 LNPs might achieve high levels of endosomal escape without inflammation.

### Endosomal escape can induce different types of endosomal damage and the endosomal damage class correlates with inflammation

In this portion of the results, we hypothesize that endosomal escape must cause some endosomal damage, including “endosomal holes.” Holes in the endosomal membrane would help mRNA reach the cytosol, but would also expose the cytosol to toxic endogenous endosomal contents, causing inflammation^42^.

We confirmed that our RNA-LNPs were localizing primarily to endosomes by visualizing co-localization of endosomes and LNPs in RAW macrophages (Fig. 3A & Supplementary Fig.11B) and A549 cells (Supplementary Fig. 11A) 30 minutes, 1h, and 6h after cKK-E12 LNP treatment. We labeled endosomes and lysosomes with Lysotracker dye and traced LNPs formulated with a fluorescent lipid (18:1 PE-TopFluor AF594). At all time points tested, there was a strong correlation between the LNP signal and the endosome/lysosome signal, as shown by the corresponding Pearson’s coefficient values (Fig. 2B). This shows that at early and late time points post-LNP administration, the lipid components of the LNPs are co-localized with the endolysosomal system (hereafter called “endosomes” for simplicity).

**Fig 3:**
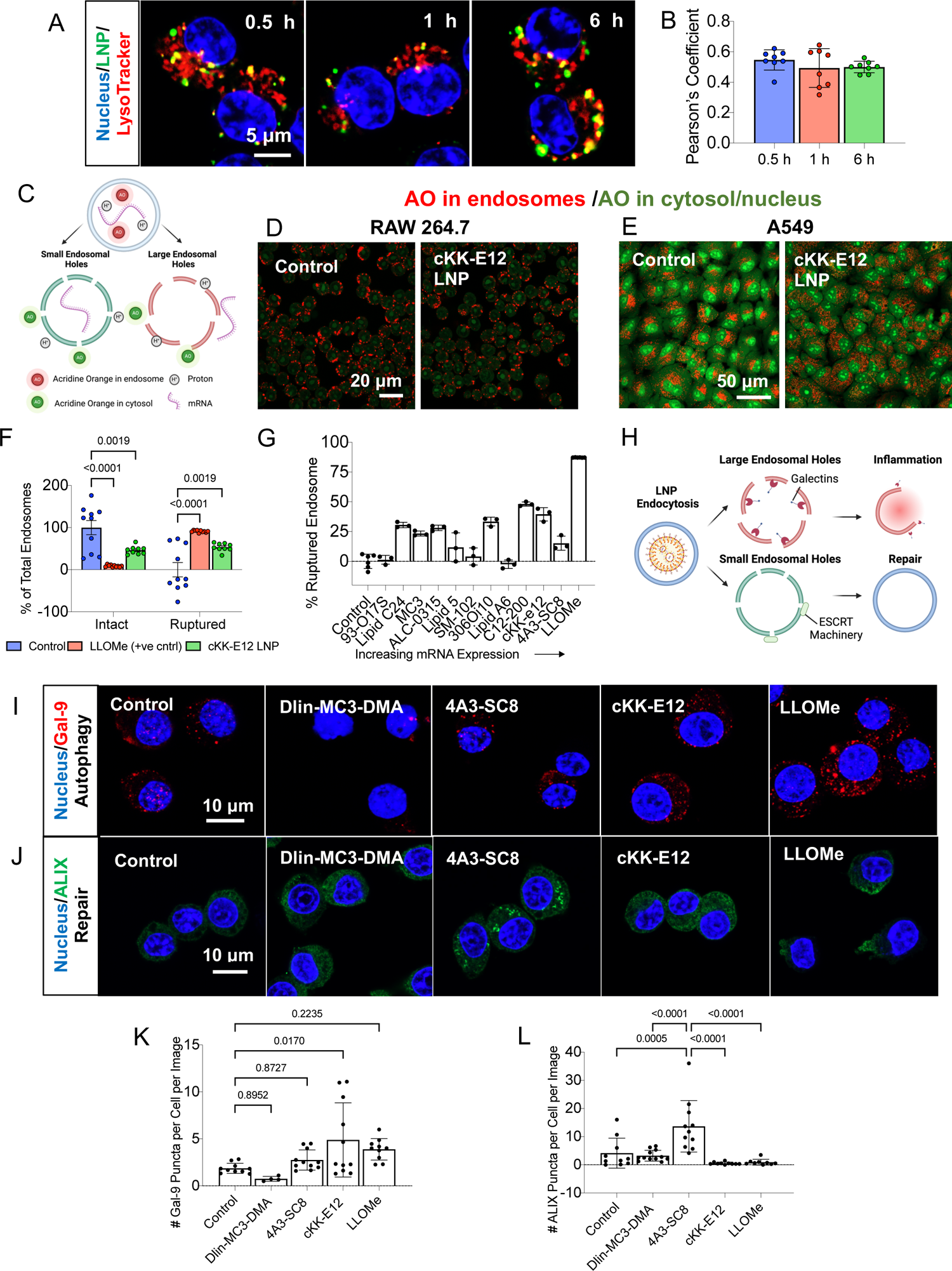
Escape of RNA payload from endosomes induces endosomal damage of different classes. (A) At 0.5h, 1h, and 6h post-treatment in RAW macrophages, LNPs are colocalized with endosomes. (B) Pearson’s coefficient values from (A). (C) Schematic showing the payload release from small and large endosomal ruptures. Small endosomal ruptures are only permeable to protons or low molecular weight molecules such as Acridine Orange (AO) while large endosomal ruptures additionally allow for the permeation of high molecular weight material such as mRNA. Control or cKK-E12 LNP-treated (D) RAW macrophages or (E) A549s were treated with AO which emits red fluorescence in acidic endosomes and green fluorescence in the nucleus or cytosol. LNP pre-treatment increases the green MFI and decreases the red MFI of AO in both cell types indicating endosomal damage. (F) Fraction of intact and ruptured endosomes in control RAW macrophages or those treated with LNPs or the endosomal rupture agent LLOME as a positive control, calculated by obtaining the ratio of the red MFI values in (C) to the green MFI values. While almost 100% of endosomes are intact in control cells, LNPs and LLOME rupture >50% and >90% of endosomes respectively. (G) Fraction of ruptured endosomes after treatment with 11 ionizable lipid LNP formulations or LLOME. (H) Schematic of the potential outcomes of damaged endosomes after LNP endocytosis. Large endosomal holes primarily recruit the sugar-binding proteins galectins which facilitate inflammatory responses. Small endosomal holes primarily recruit the ESCRT machinery which promote repair. RAW macrophages were treated with LLOME or with LNPs formulated with Dlin-MC3-dma, 4A3-SC8, or cKK-E12, and stained with (I) Galectin-9 or (J) ALIX. The number of (K) Galectin-9 or (L) ALIX puncta per cell per image from (I) and (J). cKK-E12 LNPs lead to the highest levels of galectin recruitment while 4A3-SC8 LNPs most significantly recruit the ESCRT machinery. Statistics: for (G), n=3-6. For all other graphs, n=5-12 images from 3 biological replicates. The data shown represents mean ± SEM. For all graphs, comparisons between groups were made using one-way ANOVA with Tukey’s post-hoc test.

We employed the pH-sensitive dye Acridine Orange (AO) to determine if LNPs compromise endosomes’ ability to maintain a more acidic pH than that of the cytosol. In acidic vesicles such as the endosome, AO becomes protonated, cannot leave the endosome, and fluoresces red. Under non-acidic conditions, such as in the cytoplasm or nucleus, AO fluoresces green^43^. Damage to the endosome will compromise the endosomal proton gradient, leading to leakage of AO into the cytosol, and diminishing the red fluorescence of AO. AO is a sufficiently small molecule that it should leak through holes in the endosomal membrane, even if those holes are too small to allow mRNA through (Fig. 3C). In both RAW macrophages (Fig. 3D) and A549 epithelial cells (Fig. 3E), cKK-E12 LNPs induced an increase in green AO fluorescence and a decrease in red AO fluorescence, indicating endosomal damage. We quantified the extent of endosomal damage in RAW macrophages using flow cytometry. The fraction of intact endosomes in each sample was calculated as the ratio of red AO fluorescence to green AO fluorescence, normalized to that in untreated control samples (Supplementary Fig. 12). As a positive control, we treated cells with L-leucyl-L-leucine methyl ester (LLOMe), a known lysosomotropic agent that severely damages the endosomal membrane^44^. While <1% of endosomes are ruptured in control cells, LLOME-treated cells have >90% of their endosomes ruptured (Fig. 3F). In cells treated with cKK-E12 LNPs, >50% of endosomes are ruptured. Given that it is estimated that ∼1% of cargo mRNA escape to the cytosol^45–48^, the high fraction of damaged endosomes suggests that endosomal escape of RNA comes with a large amount of collateral endosomal damage.

We performed AO endosomal rupture quantification on cells treated with LNPs formulated with some of the ionizable lipids screened in Fig. 2, 6h after LNP treatment (Fig. 3G, Supplementary Fig. 12). For select ionizable lipids, rupture was also quantified 30 minutes, 1h, 2h, and 4h after LNP treatment (Supplementary Fig. 13). Over 80% of the ionizable lipids caused endosomal rupture severe enough to detect by the AO assay. This includes ionizable lipids such as Dlin-MC3-DMA and ALC-0315 which were not significantly inflammatory and did not lead to high mRNA expression. There was thus a weak correlation between mRNA expression and AO-assessed endosomal rupture in our library of LNP formulations. As shown in Fig. 3C and noted above, AO is a small molecule (MW = 265 g/mol) compared to the mRNA used in our studies (MW >652,000 g/mol). Therefore, AO would be able to leak from the endosome through smaller holes that would be impermeable to mRNA. Endosomal escape of a larger split GFP peptide (MW = 6314g/mol), follows the trend MC3 LNPs < C12-200 LNPs < SM-102 LNPs < cKK-E12 LNPs < 4A3-SC8 LNPs, more closely matching the trend for mRNA expression (Supplementary Fig. 14). Since escape of split GFP peptide requires larger endosomal holes, this trend indicates that the amount of RNA expression and inflammation induced by LNPs is not simply a function of the *quantity* of endosomal holes, but is also determined by the *type* and *size* of endosomal holes.

To further investigate the nature of the endosomal damage induced by LNPs, we looked to cellular mechanisms for sensing different types of endosomal damage induced by pathogens, particularly by viruses, which also have to achieve endosomal escape to deliver their nucleic acid cargo^49^. Microbiology studies show that large endosomal ruptures recruit galectins, sugar-binding proteins which recognize glycans on the intra-luminal leaflet of endosomal membrane, which become exposed upon rupture. Upon detecting large endosomal holes induced by pathogens, galectins modulate inflammatory cascades and facilitate lysophagy, the process through which cells degrade damaged endosomes by fusing them with lysosomes^19,20,50^. Smaller endosomal ruptures trigger leakage of calcium ions into the cytosol, mediating the recruitment of the endosomal sorting complex required for transport (ESCRT) machinery. The ESCRT machinery consists of a group of proteins that form filaments that promote budding, leading to repair of the endosomal membrane^21–24^. The galectin and ESCRT pathways are illustrated in Fig. 3H. While ESCRT repair has been studied for different pathogens and particles, the response of the ESCRT machinery to LNP treatment is not known. In LNP treatment experiments, we therefore analyzed endosomal galectins as a marker of major endosomal damage and endosomal ESCRT proteins as a marker of reparable endosomal damage.

To assay galectin recruitment to large endosomal holes, we stained for galectins after treating RAW macrophages with Dlin-MC3-DMA, 4A3-SC8, or cKK-E12 LNPs, using LLOME as a positive control. While there are 15 galectins that we could have examined, previous research shows that staining of galectins 1, 3, 8, and 9 correlates with endosomal escape and RNA expression induced by LNPs^51–53^. We stained Gal-1, 3, 8, and 9, and quantified Gal-9 puncta 6 hours post-LNP treatment (Supplementary Fig. 15, Fig. 3I, Fig. 3K). Dlin-MC3-DMA LNPs, which cause little RNA expression and inflammation, led to the lowest levels of Gal-9 recruitment. cKK-E12 LNPs, which cause high levels of RNA expression and inflammation, led to the highest levels of Gal-9 recruitment. However, 4A3-SC8 LNPs, which induce high levels of expression and low levels of inflammation, did not significantly elevate Gal-9 puncta. Dlin-MC3-DMA and cKK-E12 LNPs were chosen as representative of the portion of our LNP library for which expression and inflammation correlate and Gal-9 puncta track with this correlation for these two formulations. But, for our outlier 4A3-SC8 LNPs, Gal-9 tracks with inflammation but not expression. This finding indicates that LNP-associated inflammation follows from the major endosomal damage marked by Gal-9, but, in the case of 4A3-SC8, galectin-associated major endosomal damage, and thus inflammation, is not necessary for endosomal escape.

To probe for lesser endosomal damage, we stained for ALG-2-interacting protein X (ALIX), a part of the ESCRT machinery^54^. We found no significant ALIX recruitment after treating RAW macrophages with Dlin-MC3-DMA LNPs, cKK-E12 LNPs, or LLOME. However, significant numbers of ALIX puncta form after treatment with 4A3-SC8 LNPs (Figs. 3J, 3L, Supplementary Fig. 16). This indicates that 4A3-SC8 LNPs cause smaller, reparable endosomal ruptures. Repair of endosomes by ESCRTs has been shown to actively limit inflammation, preventing exposure of endosomal contents to the cytosol and the inflammasome-mediated, highly inflammatory process of pyroptosis that we showed LNPs can cause in Figure 1^55,56^. 4A3-SC8 LNPs’ induction of ESCRT-associated endosomal rupture explains how these LNPs can achieve effective endosomal escape without inducing inflammation.

We hypothesize that because 4A3-SC8 has numerous esters and adjacent thioethers, it can be quickly degraded in the endosome, limiting the duration and severity of the endosomal holes it induces. We posit that 4A3-SC8’s high degree of degradability allows endosomal holes big enough for RNA escape, but still in the reparable size range. The other biodegradable lipids in our library did not induce expression as well as 4A3-SC8, probably because they do not have the same multiplicity of alkyl tails (which has previously been shown to correlate with high expression^38^), combined with esters and thioethers. Collectively, the data in Figures 2 and 3 shows that major galectin-marked endosomal damage is associated with LNP-induced inflammation, but ionizable lipids like 4A3-SC8 achieve endosomal escape without inflammation via large numbers of intermediate-sized/reparable ESCRT-marked endosomal holes.

### Inhibiting endosomal damage detection by galectins ameliorates LNP-induced inflammation

In addition to facilitating the removal of damaged endosomes through lysophagy, galectins promote inflammation by facilitating inflammasome assembly, activating and recruiting innate immune cells, and promoting cytokine function^20,57^. Having shown that LNP-induced inflammation correlates with galectin recruitment to endosomes, we hypothesized that inhibiting galectins could prevent LNP-induced initiation of inflammatory pathways moderated by galectins. We inhibited galectins using the small molecule drug thiodigalactoside (TG), which competitively binds to the carbohydrate-binding domains of galectins. We tested the efficacy of TG against inflammation induced by cKK-E12 LNPs, the most inflammatory LNPs in our library. To first verify that TG pretreatment reduces the formation of galectin puncta, we treated RAW macrophages *in vitro* with TG (2.5mg/ml) for 1 hour and then administered cKK-E12 LNPs (400ng/ml of mRNA in LNPs) for 6 hours. The data show significantly reduced numbers of Gal-9 puncta following TG pre-treatment (Fig 4A). Fitting our hypothesis, TG pre-treatment limits not just LNP-induced Gal-9 puncta, but also LNP-induced production of IL-6, TNF⍺, IL-1⍺, MCP-1 (Fig. 4B), and other pro-inflammatory cytokines (Supplementary Fig. 17).

**Fig 4:**
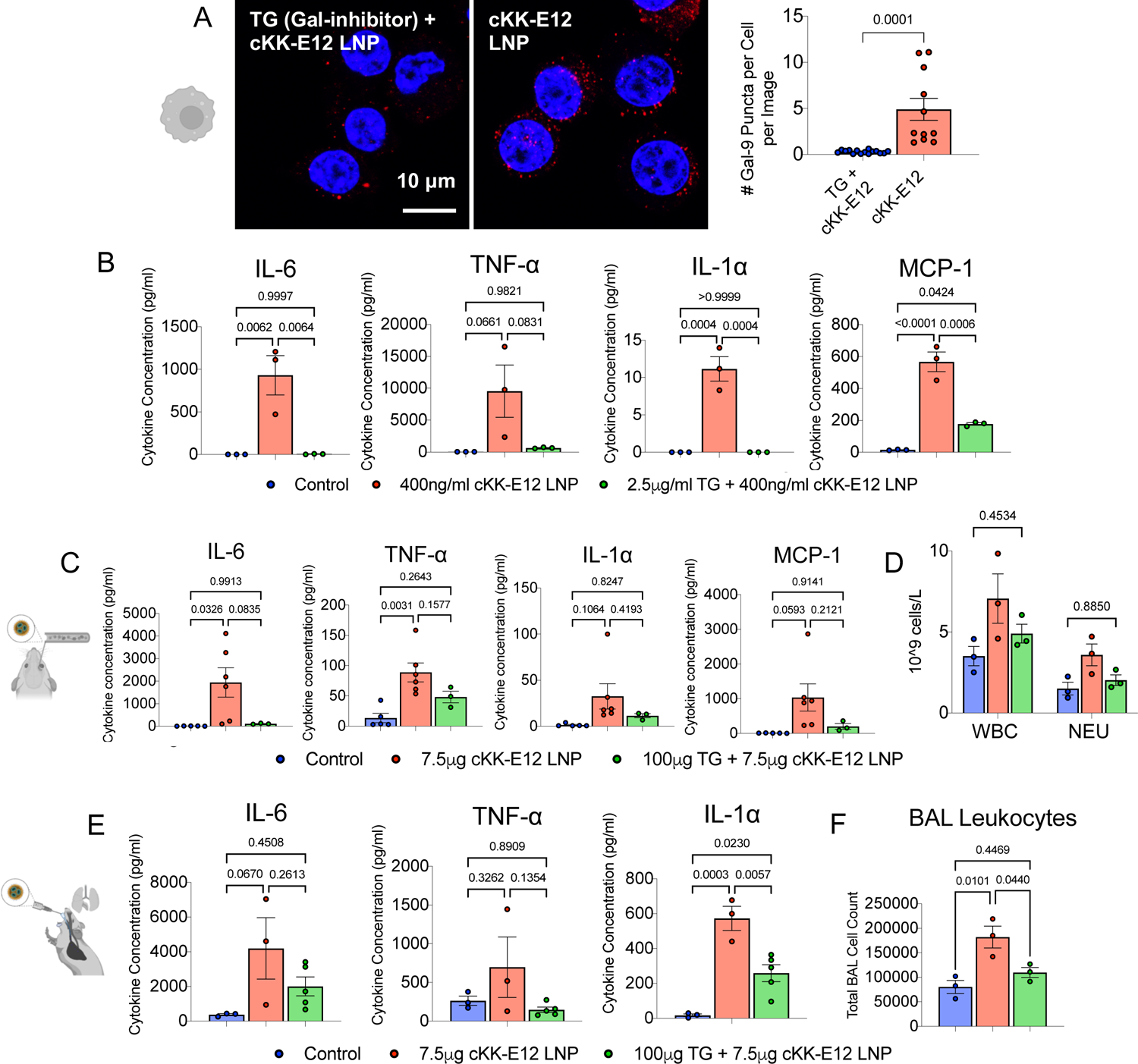
Inhibition of severe endosomal damage detection with the galectin inhibitor thiodigalactoside (TG) ameliorates LNP-induced inflammation *in vitro*, and *in vivo* across routes of delivery. (A) TG pre-treatment in RAW macrophages significantly reduces the number of galectin-9 puncta. (B) In *in vitro* RAW 264.7 macrophages, thiodigalactoside pre-treatment abrogates the LNP-induced upregulation of the pro-inflammatory cytokines IL-6, TNF⍺, IL-1⍺, and the chemokine MCP-1. (C) When TG and LNPs are administered IV into mice, TG pre-treatment significantly reduces the plasma concentrations of IL-6, TNF⍺, IL-1⍺, and MCP-1 induced by LNPs and (D) reduces the white blood cell (WBC) and neutrophil (NEU) count in the plasma to control levels. (E) When TG and LNPs are administered IT into mice, TG pre-treatment reduces the BAL concentrations of IL-6, TNF⍺, and IL-1⍺ and the (F) BAL leukocyte count. In all experiments, TG was administered 1 hour before LNP treatment. *In vitro*, cytokines were measured 6 hours after LNP administration. *In vivo,* cytokines and BAL protein and leukocyte levels were measured 2 hours after LNP treatment while blood count was measured 6 hours after LNP treatment. Statistics: for (A), n=11-15 images from 3 biological replicates. For all other graphs, n=3-6. The data shown represents mean ± SEM. For (A), comparisons between groups were made using an unpaired t-test with Welch’s correction. For all other graphs, comparisons between groups were made using one-way ANOVA with Tukey’s post-hoc test.

We also tested galectin inhibition as a preventative for *in vivo* LNP-induced inflammation. TG (100µg per mouse) was injected intravenously 1 hour before injection of cKK-E12 LNPs (7.5µg of mRNA in LNPs). Data were collected 2 hours after LNP injection. Intravenous TG pre-treatment significantly decreased LNP-induced increases in plasma levels of IL-6, TNF⍺, IL-1⍺, MCP-1 (Fig. 4C), and other pro-inflammatory cytokines (Supplementary Fig. 17). TG also prevented LNP-induced leukocytosis and neutrophilia (Fig. 4D). TG also prevented LNP-induced inflammation via the intratracheal route. TG was instilled intratracheally 1 hour before intratracheal administration of cKK-E12 LNPs. BAL fluid was extracted from the lungs 2 hours after LNP treatment. In the BAL fluid, TG significantly ablated LNP-induced increases in concentrations of IL-6, TNF⍺, IL-1⍺ (Fig. 4E), and other pro-inflammatory cytokines (Supplementary Fig. 17). Furthermore, TG reduces the LNP-induced increase in leukocyte infiltration into the alveolar space, as measured by BAL leukocyte counts (Fig. 4F). TG reduction in BAL leukocyte count persists even 24 hours after LNP instillation (Supplementary Fig. 17C). Notably, TG prevents the inflammation induced by other ionizable lipids (Supplementary Fig. 17B). Thus, galectin inhibition prevents LNP-induced inflammation *in vitro* and across multiple routes of *in vivo* LNP delivery.

### Inhibition of endosomal escape detection positively impacts mRNA expression

After showing that galectin inhibition ameliorates LNP-induced inflammation, we sought to determine its effects on RNA expression. We treated RAW macrophages with TG (2.5mg/ml) for 1 hour, administered cKK-E12 LNPs formulated with luciferase mRNA, and then measured luciferase expression 6 hours later. TG pre-treatment increases mRNA expression >2.5-fold over cKK-E12 LNPs alone (Fig. 5A). *In vivo*, intratracheal TG (100µg per mouse) given 1 hour before intratracheal cKK-E12 LNPs has no effect on luciferase expression at 6 hours after LNP treatment (Fig. 5B). However, intravenous TG (100µg per mouse) leads to ∼2.7-fold and ∼2.4-fold increases in cKK-E12 LNP-induced luciferase expression in the liver and spleen, respectively (Fig. 5C). These results indicate that in addition to ameliorating inflammation induced by LNPs, inhibition of major endosomal damage detection also generally improves RNA expression.

**Fig 5:**
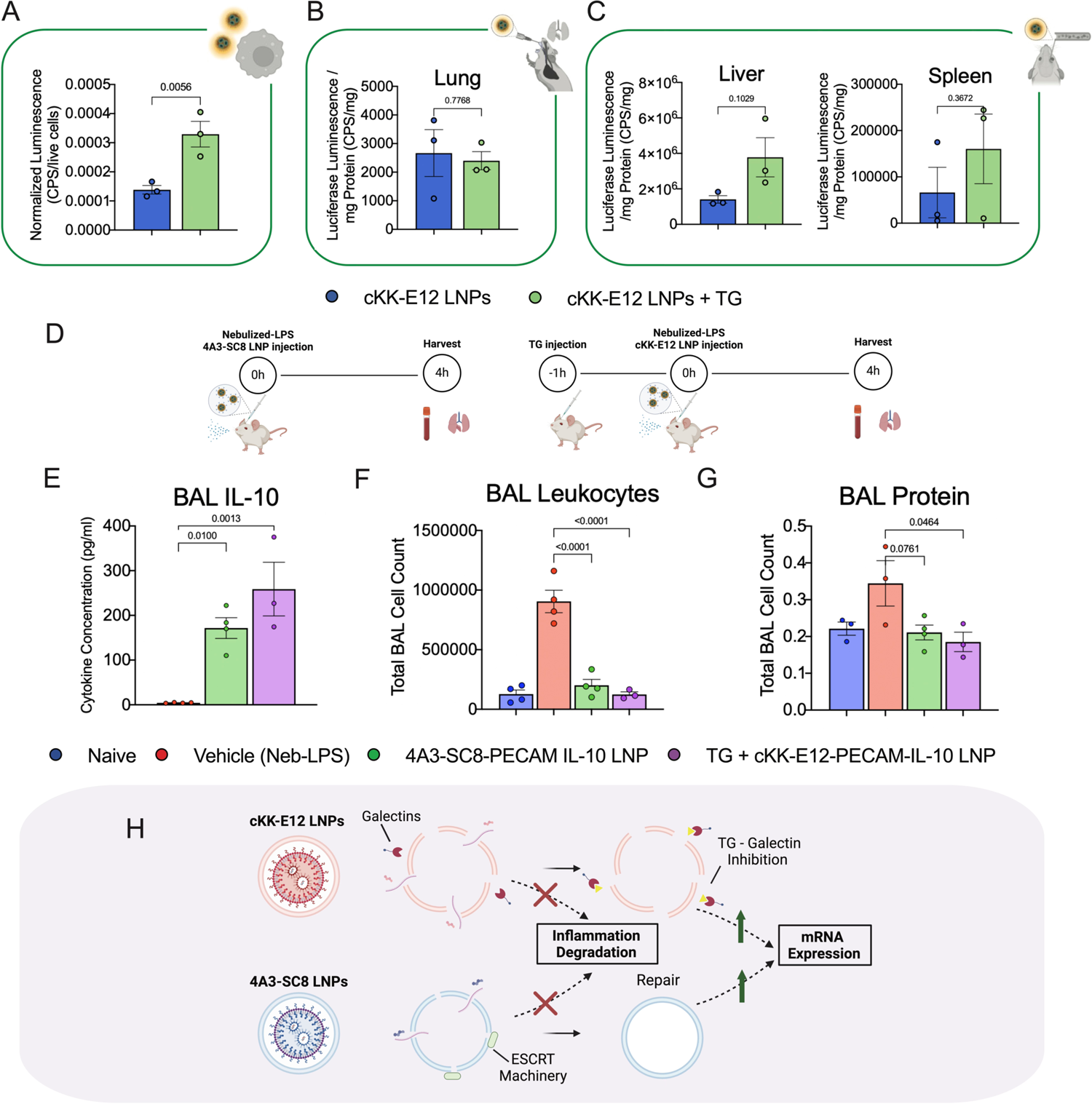
Inhibition of endosomal escape detection has positive effects on mRNA expression and non-inflammatory LNPs abrogate ARDS. (A) In RAW 264.7 macrophages, TG pre-treatment increases luciferase mRNA expression by > 2-fold. (B) TG pre-treatment does not attenuate expression in the lung when TG and LNPs are administered IT. TG pre-treatment improves mRNA expression in the (C) liver and spleen by ∼2.7 and ∼2.4-fold respectively when TG and LNPs are injected IV. In all experiments, TG was administered 1 hour before LNPs, and luminescence measurements were taken 6 hours after LNP treatment. (D) Timeline of treatments in nebulized-LPS ARDS model with LNPs containing mRNA encoding for the anti-inflammatory cytokine IL-10. (E) IL-10 mRNA-loaded 4A3-SC8 LNPs and TG + IL-10 mRNA-loaded cKK-E12 LNPs induced upregulation of IL-10 by ∼40-fold and ∼60-fold respectively in the BAL fluid. IL-10 4A3-SC8 LNPs and TG + IL-10 cKK-E12 LNPs completely abrogated (F) leukocyte infiltration (BAL leukocyte count) and (G) capillary leak into the alveolar space (BAL protein levels). (H) Summary schematic showing the two strategies for ameliorating LNP-induced inflammation while maintaining high mRNA expression namely utilizing ESCRT-recruiting ionizable lipids and inhibiting galectins. Statistics: n=3–4 and the data shown represents mean ± SEM; For (A) - (C), comparisons between groups were made using an unpaired t-test with Welch’s correction. For all other graphs, comparisons between groups were made using one-way ANOVA with Tukey’s post-hoc test.

### Non-inflammatory LNPs loaded with therapeutic mRNA cargo ameliorate ARDS

Our results demonstrate amelioration of LNP-induced inflammation while preserving high mRNA expression using; 1) an ESCRT-recruiting ionizable lipid; 2) galectin inhibition. Below, we use both of these strategies to achieve therapeutic effects with LNPs in a model of acute inflammatory lung injury. We and others have shown that LNPs exacerbate pre-existing inflammation^6^, making use of inflammatory LNPs prohibitive in patients with conditions including ARDS, stroke, and heart attack. We tested our non-inflammatory LNP formulations in a mouse model of ARDS-like acute lung injury induced by administering nebulized LPS. Despite the high mortality of ARDS patients (>45%), particularly in cases associated with COVID-19^58^, there are currently no mRNA LNP therapeutics being tested for this disease. Because LNPs severely increase inflammation in the nebulized-LPS ARDS model, preclinical testing of LNPs for ARDS treatment is impeded (Supplementary Fig. 18). We attempted to ameliorate LNP-induced inflammation in this model using small molecule drugs (dexamethasone, MCC950), proteins (IL-1Ra), and antibodies (anti-IL-6), but none of these therapeutic approaches were efficacious (Supplementary Fig. 19). We also attempted to understand the signaling pathways underlying LNP-induced inflammation exacerbation using knockout mice (ASC, CCR2, MyD88, caspase-3) but were unable to isolate a clear pathway (Supplementary Fig. 20).

Implementing our new strategies for reducing LNP-induced inflammation, we tested; 1) 4A3-SC8 LNPs; and 2) cKK-E12 LNPs with TG pre-treatment as agents for delivery of mRNA encoding the anti-inflammatory cytokine IL-10, which has been suggested as a potential ARDS therapeutic^59^, in the nebulized LPS mouse model of ARDS. We targeted LNPs to the lung by conjugating them to antibodies against PECAM (Platelet Endothelial Cell Adhesion Molecule)^60^. Mice were either intravenously injected with TG (100µg per mouse) followed by cKK-E12 LNPs 1 hour later (7.5µg of mRNA in LNPs) or intravenously injected with 4A3-SC8 LNPs (7.5µg of mRNA in LNPs). Nebulized LPS was administered immediately following LNP injection and mice were sacrificed 4 hours later (Fig 5D). 4A3-SC8 and TG + cKK-E12 LNPs induced potent upregulation of IL-10 in the BAL fluid, by ∼40-fold and ∼60-fold, respectively (Fig. 5E). Both formulations also reduced the BAL concentrations of pro-inflammatory cytokines such as IL-6 and TNF-⍺, compared to untreated levels (Supplementary Fig. 21). Most significantly, IL-10 4A3-SC8 LNPs and TG + IL-10 cKK-E12 LNPs *completely abrogated* the hallmark physiological characteristics of ARDS: leukocyte infiltration (Fig. 5F) and capillary leak (Fig. 5G) into the alveolar space, even at longer timepoints (Supplementary Fig. 22). TG pre-treatment alone did not affect ARDS phenotypes (Supplementary Fig. 23).

We have demonstrated that we can ameliorate LNP-induced inflammation by; 1) inhibiting galectins, thus preventing sensing of severe, irreparable endosomal damage and eliminating downstream galectin-mediated inflammatory responses; 2) formulating LNPs with branched, multi-tail, highly biodegradable ionizable lipids such as 4A3-SC8 which induce less severe, reparable endosomal damage that can be repaired by ESCRT proteins before inflammation is triggered (Fig. 5H).

## Discussion

Despite the success of the mRNA-LNP COVID vaccines, LNP-induced inflammation presents one of the greatest challenges in the field, preventing the translation of LNPs for other indications, especially those with pre-existing inflammation. While several hypotheses have been extended, the mechanisms behind LNP-induced inflammation remain poorly understood^10^. In this study, we demonstrate that endosomal escape of RNA from LNPs, which is necessary for expression, also induces endosomal damage that triggers inflammation.

Through structure-function screening of various ionizable lipids, we show that there is a positive correlation between RNA expression and inflammation. We demonstrate that the highest-expressing, most inflammatory ionizable lipids also tend to induce the most severe endosomal damage. If uncontrolled, endosomal damage and exposure of the endosomal lumen can cause leakage of harmful substances such as cathepsins, protons, and reactive oxygen species, triggering inflammatory pathways like the inflammasome^7^. However, our screening revealed an outlier ionizable lipid, 4A3-SC8, that induces high RNA expression without causing significant inflammation.

Mechanistic investigation of cell responses to endosomal damage revealed that LNP-induced inflammation is dependent on both the quantity of damaged endosomes and the size of endosomal holes generated. Large endosomal holes are sensed by galectins, which detect exposure of glycans on the inner leaflet of the endosome. If endosomal injury is too severe, galectins modulate autophagy, where the autophagosome degrades the endosomal membrane remnants^19,20,50^. However, smaller holes in the endosome recruit the ESCRT machinery to repair the damaged membrane^21,22,50^. ESCRT processes may limit inflammation by preventing exposure of inflammatory endosomal contents to the cytosol. We show that LNPs formulated with 4A3-SC8 cause recruitment of ESCRT machinery to endosomes, suggesting that 4A3-SC8 causes enough endosomal membrane damage to facilitate high RNA expression, but the damage is repaired by ESCRT proteins. In our screening library, 4A3-SC8 LNPs were unique in causing both high RNA expression and low inflammation. Among our other tested LNPs, low inflammation could only be achieved when RNA expression was sacrificed.

4A3-SC8 was also unique among our tested ionizable lipids for having *both* biodegradable ester and thioether linkages and a multi-tail structure that is better able to form the inverse hexagonal phase that has been linked to more efficient endosomal escape. Specifically, thioether groups can undergo redox-triggered degradation^61,62^. We therefore posit that the structure of 4A3-SC8 may stimulate frequent endosomal membrane damage, but also rapid degradation and clearance of the offending lipid, allowing rapid ESCRT-mediated repair before the endosomal damage leads to irreparable rupture. This study is the first to report ESCRT recruitment by LNPs and link the amelioration of LNP-induced inflammation to ESCRT repair. This provides a paradigm for designing ionizable lipids that are high-expressing and non-inflammatory.

In our studies, prophylactic galectin inhibition ameliorates LNP-induced inflammation *in vitro* and *in vivo*. The pan-galectin inhibitor thiodigalactoside (TG) also generally increases mRNA expression. Galectins play a role in sensing severe endosomal damage, including that induced by LNPs. Our results demonstrate for the first time that galectins have a net pro-inflammatory effect following LNP-induced endosomal damage. This may be due to several mechanisms that warrant further investigation. Galectins have been shown to promote NLRP3 inflammasome activation, particularly in macrophages^63,64^. Our data is consistent with the link between LNP endosomal escape, galectins, and inflammasome activation: Galectin inhibition reduced the concentration of IL-1β (in cases where this cytokine is upregulated by LNPs), which is produced by the NLRP3 inflammasome (Supplementary Fig. 17). Noting that galectins promote leukocyte recruitment by modulating immune cell adhesion or chemoattraction^65–70^, we have demonstrated that intratracheal galectin inhibition ameliorates LNP-provoked leukocyte infiltration into the alveolar space (Fig. 4F) and intravenous galectin inhibition reduces LNP-provoked increased in circulating leukocytes (Fig. 4D).

Our investigation of the signaling pathways underlying LNP-induced inflammation resulted in LNP formulations that efficiently deliver RNA without exacerbating existing inflammation, in turn leading to an RNA therapeutic for ARDS. The principles behind the development of these non-inflammatory LNP formulations can be harnessed to expand the therapeutic potential of RNA-LNPs and can be applied to other inflammatory conditions such as heart attack and stroke.

## Materials and Methods

### Materials

DOPE (1,2-dioleoyl-sn-glycero-3-phosphoethanolamine), cholesterol, DMG-PEG 2000 (1,2-dimyristoyl-rac-glycero-3-methoxypolyethylene glycol-2000), 18:1 PE TopFluor AF 594 (1,2-dioleoyl-sn-glycero-3-phosphoethanolamine-N-(TopFluor® AF594) (ammonium salt)), and DSPE-PEG2000-azide (1,2-distearoyl-sn-glycero-3-phosphoethanolamine-N-[azido(polyethylene glycol)-2000] (ammonium salt)) were purchased from Avanti Polar Lipids. Ionizable lipids 4A3-SC8, cKK-E12, SM-102, ALC-0315, and C12–200 were purchased from Echelon Biosciences. All other ionizable lipids were purchased from Broadpharm. L-leucyl-L-leucine, methyl ester (LLOMe) was purchased from Cayman Chemical. Thiodigalactoside was purchased from MedChemExpress.

### Animals

All experiments involving animals were conducted following the guidelines outlined in the Guide for the Care and Use of Laboratory Animals (National Institutes of Health, Bethesda, MD). Approval for all animal protocols was obtained from the University of Pennsylvania Institutional Animal Care and Use Committee. Male C57BL/6 mice aged 6–8 weeks (23-25 g) sourced from The Jackson Laboratory, Bar Harbor, ME, were utilized for the experiments. The mice were housed in a controlled environment at temperatures between 22–26°C, adhering to a 12/12h dark/light cycle, and had unrestricted access to food and water.

### LNP Formulation

LNPs were formulated using the microfluidic mixing method. An organic phase containing a mixture of lipids dissolved in ethanol at a designated molar ratio (Supplementary Table 1) was mixed with an aqueous phase (50 mM citrate buffer, pH 4) containing 5moU modified Luciferase mRNA (unless otherwise stated) that was purchased from TriLink Biotechnologies, at a flow rate ratio of 1:3 and at a total lipid to mRNA weight:weight ratio of 40:1 in a microfluidic mixing device (NanoAssemblr Ignite, Precision Nanosystems). LNPs were dialyzed against 1× PBS in a 10 kDa molecular weight cut-off cassette for 2 h, sterilized through a 0.22 μm filter, and stored at 4 °C. IL-10 mRNA was synthesized via *in vitro* transcription as described previously^6^.

### LNP Characterization

Measurements of hydrodynamic nanoparticle size, distribution, polydispersity index, and zeta potential were conducted through dynamic light scattering using a Zetasizer Pro ZS from Malvern Panalytical. The encapsulation efficiencies and concentrations of LNP RNA were determined using a Quant-iT RiboGreen RNA assay (Invitrogen).

### Cell Culture

Raw264.7 mouse macrophages were purchased from ATCC and cultured in Dulbecco’s modified Eagle’s medium (DMEM) with 10% heat-inactivated fetal bovine serum (FBS) and 1% penicillin/streptomycin (PS). A549_GFP1-10_ cells were generously gifted by Prof. Andrew Tsourkas and cultured in DMEM with 10% FBS, 1% PS and 2 ug/mL puromycin. MLE-12 lung epithelial cells were kindly gifted by Prof. Jeremy Katzen and cultured in DMEM-F12K containing 2% FBS, 1% PS, 10 nM β-estradiol, 10 nM hydrocortisone, 2 mM L-glutamine and 1:100 insulin-transferrin-selenium-ethanolamine supplement. All cells were incubated with 5% CO_2_ at 37 ℃.

For 24 well-plate experiments, RAW 264.7 macrophages were seeded at 400k density using 500 uL volume and treated with 300 uL of LNP volume (diluted in DMEM). A549_GFP1-10_ and MLE-12 cells were seeded at a density of 60k.

### *In Vitro* Luciferase Delivery for measurements of mRNA Transfection Efficiency

Luciferase mRNA LNPs fabricated as described above were incubated with cells for 6 hours. Supernatants were collected (spun down at 10,000 x g for 15 mins) for cytokine analysis using LegendPlex 13-plex Mouse Inflammation Panel (Biolegend). Adhered cells were washed with 1x PBS and collected using 0.25% trypsin. Cell counts were determined using Cell Countess after staining with trypan blue to determine cell viability. Then, cells were spun down at 300 x g for 5 mins to pellet, washed with 1x PBS, and lysed with 120 uL of 1x Promega Luciferase Assay System Cell Culture Lysis Reagent. For luciferase expression, 20 µL of lysed sample was loaded onto a white 96 well-plate, and the luminescence was read on a luminometer (Promega) after 100 µL of luciferin solution (Promega) was added well by well by an autosampler. Final luminescence readings were then normalized based on cell count.

### *In Vivo* Luciferase Delivery for measurement of mRNA Transfection Efficiency

Luciferase mRNA LNPs fabricated as described above were injected into mice intravenously or instilled intratracheally for a circulation time of 6 hours. Select organs were then flash-frozen until the day of analysis or homogenized immediately. Samples were suspended in 900 uL of homogenization buffer (5mM EDTA, 10mM EDTA, 1:100 diluted stock protease inhibitor (Sigma), and 1x (PBS), samples were then loaded with a steel bead (Qiagen), then placed in a tissue homogenizer (Powerlyzer 24, Qiagen) using the following settings: Speed (S) 2000 rpm, 2 Cycles (C), T time 45 sec, and pause for 30 sec). After this, 100 uL of lysis buffer (10% Triton-X 100 and PBS) was added into each tube and then allowed to incubate for 1 hr at 4C. After this, they were immediately transferred into fresh tubes, and sonicated, using a point sonicator to remove in excess DNA, using an amplitude of 30%, 5 cycles of 3 secs on/off. After this, samples were then centrifuged at 16,000 x g for 10 minutes. The resultant lysate was either frozen or prepared for luminometry analysis.

For luciferase expression, 20 µL of undiluted sample was loaded onto a white 96 well-plate and the luminescence was read on a luminometer (Promega) after 100 µL of luciferin solution (Promega) was added well by well by an autosampler. Last, a DC Protein Assay (Bio-Rad) was performed according to manufacturer specifications using diluted samples, specifically a 1:40 dilution for lung and spleen tissues and a 1:80 dilution for liver tissues. Final luminescence readings were then normalized based on total protein concentration obtained from Lowry Assay.

### Bronchoalveolar lavage (BAL) Protein and Leukocyte measurements

mRNA LNPs were administered at various doses for various time points. After blood collection, bronchoalveolar lavage was performed by inserting a catheter intratracheally, dispensing 800uL of ice cold 1x PBS (0.5uM EDTA) using 1mL BD syringe and performing a total of three washes prior to collection. Total leukocyte count was measured by diluting BAL fluid with trypan blue (1:1) and using cell countess for quantification. BAL fluid was spun down at 300g for 5 mins to pellet cells. The supernatant was collected and used to perform DC Protein Assay (Bio-Rad) using manufacturer instructions for total BAL protein quantification. Total BAL protein and leukocyte count was normalized using total BAL volume collected.

### Cell viability assay

Cells were seeded in 96-well plates and incubated overnight (Raw264.7 cells: 1E5 cells/well; MLE-12 cells: 8E3 cells/well). Following different LNP treatments, 10% cell counting kit-8 (CCK-8, ALX850039KI01, Enzo Life Sciences) reagent in complete medium was added to cells for 2 h incubation. Then, CCK-8 absorbance was recorded by microplate reader at 450 nm (660 nm absorbance as reference). Cell viability rates were normalized to control cells.

### Apoptosis/Necrosis/Pyroptosis Assay

Apoptotic cells were detected with fluorescent phosphatidylserine. Necrotic cells were detected with fluorescent 7-AAD. Pyroptotic cells were detected with fluorescent caspase-1. Healthy cells were detected with fluorescent cytocalcein. Markers were then quantified with flow cytometry. All markers were purchased from Abcam and used according to the manufacturer’s instructions.

### Complete Blood Count (CBC)

Blood was collected from mice drawing from the inferior vena cava (terminal procedure). VetScan was used to obtain cell counts from whole blood according to the manufacturer’s instructions.

### *In Vitro* and In Vivo Cytokine Measurements

Cytokine measurements were carried out on plasma, BAL, or cell culture supernatant with a LegendPlex 13-plex Mouse Inflammation Panel (Biolegend) according to the manufacturer’s instructions.

### Pig Lung Cytokine Measurements

To inflate *ex vivo* pig lungs, a 6.0 ET tube was cannulated into a lobe with least damage. The ET tube was secured into the airway using surgical ties, once the tube was secured, the ET cuff was inflated. A four-way connector was used to attach air flow applying continuous flow around 2-3L/min, and tubing to a sealed Erlenmeyer flask with a side arm to maintain 5cm H_2_O positive end-expiratory pressure (PEEP). The top of the connector was sealed using parafilm to maintain an appropriate level of inflation without damaging the lung tissue. Before instilling solution, the lungs were perfused. To instill LNP solution, another ET tube was used and secured in the airway with surgical ties as done previously. For control groups, around 30 cc of 1xPBS was installed slowly via an 8F catheter snaked through the ET tube. The lobe was massaged during installation. The ET tube was then taken out, adjusted to deliver to a different lobe, and secured using surgical ties. This process was repeated in the other lobe to instill around 20 cc of LNP solution (0.8mg of mRNA in LNPs). Once complete, the catheter was removed and the end of the ET tube was sealed with parafilm. To ensure little leakage of air and solution, both ET tubes were tied again with surgical ties. Keeping both ET tubes upwards, the lung was placed into a ziplock bag and incubated in a water bath at 37℃ for 3 hours. BAL was then harvested. TNF-⍺ and IL-6 pig ELISAs were purchased from Abcam and used according to the manufacturer’s instructions to measure BAL cytokine levels.

### hMDM Cell Culture and isolation

Human peripheral blood mononuclear cells (PBMC) were separated from blood obtained from de-identified healthy donors (New York Blood Center, Long Island City, New York) by Ficoll-Paque (GE Healthcare, Piscataway, NJ, USA) gradient centrifugation. Following isolation, the percentage of monocytes in the PBMC was quantified using a monocyte isolation kit (MACS, Miltenyi Biotechnology) and PBMC were plated at a density of approximately 1 x 105 monocytes/cm2 to obtain a pure culture of human monocyte-derived macrophages (hMDM) via adherence isolation. PBMC were cultured in RPMI-1640 supplemented with 10% FBS, 5% human AB serum, 10 mM HEPES, 1% penicillin/streptomycin, and M-CSF (10 ng/mL) for 3 days then washed with fresh media to remove non-adherent cells. Adherent cells were cultured another 3 days in fresh media containing M-CSF. After 6 days in culture, cells are considered matured to hMDM. Experiments were performed at day 6 or 7.

### hMDM Immunocytochemical Staining and Analysis

Human monocyte-derived macrophages (hMDM) were cultured in Nunc™ MicroWell™ 96-well optical-bottom plates (Thermo Fisher Scientific, Waltham, MA) and left untreated or treated with vehicle (sterile 1X PBS), LPS (1ng/mL) or LNPs for 90 minutes. Treatment with LPS (1 ng/mL) for 90min was used as a positive control. Cells were then fixed using 4% paraformaldehyde for 10 mins and permeabilized with 0.1% Triton X-100 in PBS for 5 min. Cells were blocked for 30 min in 1% BSA and 300 mM glycine in 0.1% Tween-20 in PBS. The primary antibody rabbit monoclonal NF-κB (CST8242, 1:400) diluted in blocking solution and incubated at 4 C overnight. Alexa Fluor 488 goat anti-rabbit secondary antibody (Thermo Fisher, 1:1000) was used for detection, nuclei were stained with DAPI (0.2 μg/mL) and plasma membranes were stained with cell mask deep red (CMDR, Thermo Fisher Scientific, 250 ng/mL). Images were acquired on a Cell Insight CX7 High Content screening platform, an automated confocal scanning microscope, acquiring 20 fields per well using a 20x objective. Four wells were imaged for each condition, acquiring approximately 1,000 - 1,500 cells per well (4,000 – 6,000 cells per condition). Images were acquired with a fixed exposure time of 0.07 s, 1.68 s, 0.035 s (DAPI, NF-κB, CMDR, respectively) and intra-well autofocusing at every field. Additional paramaters used for these analyses are found in Supplementary Table 2. Images were analyzed using HCS studio software and the Cellomics Colocalization bio-application (Cellomics, ThermoFisher, Pittsburgh, PA). This analysis creates a binary mask for the cytoplasmic (cell mask deep red) and nuclear (DAPI) regions of interest (ROI), and then quantifies the intensity of NF-κB staining in each ROI for every cell. The NF-κB intensity in the nuclear ROI is divided by the NF-κB intensity in the cytoplasmic ROI in each individual cell to generate a nuclear colocalization ratio for each individual macrophage. This quantifies the relative amount of NF-κB in the nucleus, while also controlling for cell size and differences in the total amount of NF-κB in different cells. The nuclear colocalization ratio is averaged across all cells and wells from a particular condition, and then the average ratios from all conditions across all donors are compared using an ANOVA.

### Acridine orange assay by confocal imaging and flow cytometer

Lysosomal pH changes were tested by acridine orange staining. To analyze acridine orange by flow cytometry, Raw264.7 cells were seeded at 4e5 cells/well and then incubated with 400 ng/mL LNP for 6 h. For positive control, 6.8 mM LLOMe was cocultured with cells for 1 h. Then, cell pellets were harvested and stained by 1 ug/mL acridine orange for 10 min. At the end of staining, add 1 mL/tube PBS immediately. Then cells were washed by centrifugation at 4℃, 300 g, 5 min and finally analyzed by Guava easyCyte flow cytometer (Luminex). To record acridine orange images, cells were seeded to 8-well μ-Slide chambers (Raw264.7 cells: 1.5E5 cells/well; A549_GFP1-10_ cells: 1.5E4 cells/well) and then treated with LNP for 6 h. As for the positive control group, Raw264.7 cells were incubated with 6.8 mM LLOMe for 1 h and A549_GFP1-10_ cells were treated with 68 μM LLOMe for 2 h. Next, cells were labeled by 10 μg/mL acridine orange for 10 min and then imaged by LSM980 confocal microscope (Zeiss).

### Colocalization imaging of LNPs and endo-lysosomes

Cells were seeded to 8-well μ-Slide chambers (Raw264.7 cells: 1.5e5 cells/well; A549_GFP1-10_ cells: 1.8e4 cells/well) and then treated with cKK-E12 LNPs formulated with 0.3 mol % of 18:1 TopFluor PE-Alexa Fluor 594 for 0.5, 1 and 6 h. Endo-lysosomes were labeled by 200 nM LysoTracker DeepRed (L12492, Invitrogen) for 30 min and nuclei were stained by Hoechst33342 (R37650, Invitrogen). Fluorescent images were acquired by LSM980 microscopy (Zeiss) and colocalization of LNPs and endo-lysosomes was analyzed by Pearson’s coefficient in ImageJ.

### Split GFP assay

A549_GFP1-10_ cells were seeded at 5.5e4 cells/well in 24-well plates. As reported previously^71^, 2 µL of Lipofectamine (13778030, Thermo Fisher) was firstly mixed with 8 uL Opti-MEM, and then 2.87 µL 1mg/mL S11 protein was added by gender pipetting. The final mixture was incubated at room temperature for 15 min. Next, the Lipofectamine:S11 mixture was added to cells as the total S11 concentration of 500 nM, followed by a 6-h incubation. For the LLOMe-treated group, cells were pretreated with 3.4 mM LLOMe for 30 min and then cocultured with S11 protein or lipofectamine:S11 mixture for 6 h. For LNP delivery, S11-loaded LNPs were diluted with complete medium and then added to cells for 6 h. Split GFP expression was measured by Guava easyCyte flow cytometer (Luminex). S11 peptides were synthesized (Lifetein) as reported previously^71^.

### Galectin and Alix puncta imaging and quantification

After LNP and LLOMe treatments, cells were fixed in 4% fresh paraformaldehyde for 15 min. Permeability was then performed with 0.05% saponin buffer (J63209.AK, Invitrogen) for 10 min. Then, samples were incubated with 10% normal goat serum (50062Z, Life Technologies Corp.) for 1 h. After PBS washing, cells were cocultured with 1:150 primary antibodies (anti Gal-1, sc-166618, Santa Cruz Biotechnology Inc.; anti-Gal-3, sc-32790, Santa Cruz Biotechnology Inc; anti-Gal-8, sc-377133, Santa Cruz Biotechnology Inc; anti-Gal-9, ab275877, Abcam; anti-Alix, ab275377, Abcam) at 4℃ overnight. Unbound antibodies were washed with PBS. Next, 1:750 secondary antibodies were added to cells (Alexa Fluor 488-conjugated goat anti-rabbit antibody, A11008; Alexa Fluor 647-conjugated goat anti-rabbit antibody, A21244; Alexa Fluor 647-conjugated goat anti-mouse antibody, A11029; Alexa Fluor 488-conjugated goat anti-mouse antibody, A11008; all from Invitrogen) and incubated at 37 ℃ for 1.5 h. Cell nuclei were labeled by DAPI. Images were acquired by LSM980 microscopy (Zeiss).

For intracellular puncta quantification, images were analyzed with Gaussian blur function by ImageJ. Simply, images were duplicated and processed with Gaussian blur to obscure dispersed background signals. Then, image subtraction was performed to remove the background noises. For Gal-9 puncta, the radius sigma was set as 1 for the duplicated image. Intracellular Gal-9 puncta were selected by size (>5). For Alix puncta, the raw image was set blurring with the radius sigma of 1 while the duplicated one was performed by the radius sigma of 2. Alix puncta were selected by size (>10). Total puncta numbers were measured and normalized to cell number in each image.

### Nebulized LPS Model

Mice were exposed to nebulized LPS in a “whole-body” exposure chamber, with separate compartments for each mouse (MPC-3 AERO; Braintree Scientific, Inc.; Braintree MA). To maintain adequate hydration, mice were injected with 1 mL of sterile saline, 37°C, intraperitoneally, immediately before exposure to LPS. LPS (L2630-100 mg, Sigma Aldrich) was reconstituted in PBS to 10 mg mL−1 and stored at −80 °C until use. Immediately before nebulization, LPS was thawed and diluted to 5 mg mL−1 with PBS. LPS was aerosolized via a mesh nebulizer (Aerogen, Kent Scientific) connected to the exposure chamber (NEB-MED H, Braintree Scientific, Inc.). 5 mL of 5 mg mL−1 LPS was used to induce the injury. Nebulization was performed until all liquid was nebulized (≈20 min).

### Statistics

All results are expressed as mean ± SEM unless specified otherwise. Statistical analyses were performed using GraphPad Prism 8 (GraphPad Software) * denotes p<0.05, ** denotes p<0.01, *** denotes p<0.001, **** denotes p<0.0001.

## Acknowledgments

Research reported in this publication was supported by the American Heart Association under Grant 23PRE1014444 (to S.O.-L.), NIH F31 fellowship (award number 1F31AG077874-01) (to M.L.A), Ruth L. Kirschstein National Research Service Award (NRSA) F31HL154662 (to M.E.Z.), Grant NIH R61DA058501, R01DA057337 (to P.J.G.) and Grant NIH R01 HL157189 (to V.M., J.W.M., and J.S.B).

