## Supplementary Figures and Captions for "Lipid Nanoparticle-Associated Inflammation is Triggered by Sensing of Endosomal Damage: Engineering Endosomal Escape Without Side Effects"

| Ionizable Lipid | Size (nm) | PDI |
| --- | --- | --- |
| 🟢 93-017S | 102.8 | 0.097 |
| 🔴 Lipid C24 | 114.5 | 0.069 |
| 🟡 DLin-MC3-DMA | 80.97 | 0.226 |
| 🟣 🔴 1O14 | 281.6 | 0.269 |
| 🟢🔴 ALC-0315 | 78.68 | 0.156 |
| 🟢 🔴 Lipid 5 | 88.43 | 0.207 |
| 🟣 🟢🔴 Lipid AX4 | 136.1 | 0.077 |
| 🟢🔴 SM-102 | 72.32 | 0.178 |
| 🟣🟢🔴 306Oi10 | 128.7 | 0.028 |
| 🟢 Lipid A6 | 108.7 | 0.097 |
| 🟣🟢 BAMEA-O16B | 127.3 | 0.193 |
| 🟣 🔴 C12-200 | 158.1 | 0.121 |
| 🟣 🔴 98N12-5 | 115.6 | 0.209 |
| 🟣 🔴 cKK-E12 | 132.2 | 0.15 |
| 🟣🟢🔴 4A3-SC8 | 124.4 | 0.073 |

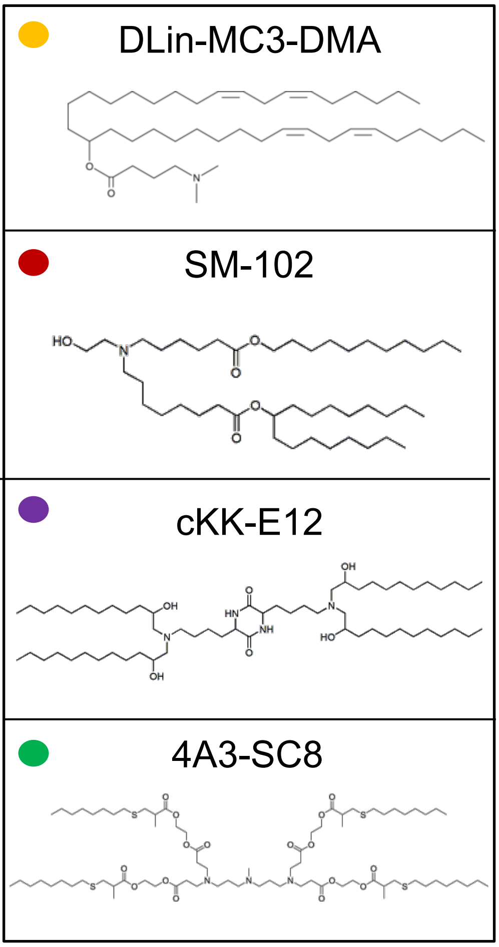


🟣 Multi tail 🔴 Branched tail

🟢 Biodegradable 🟡 Unsaturated

**Supplementary Table 1 | Ionizable lipid classification and LNP Size and PDI.** All LNP formulations consisted of 50% ionizable lipid, 38.5% cholesterol, 10% DOPE, and 1.5% DMG-PEG2K. Ionizable lipids are categorized into 4 groups indicated by colored dots. Representative structures are also shown on the right.

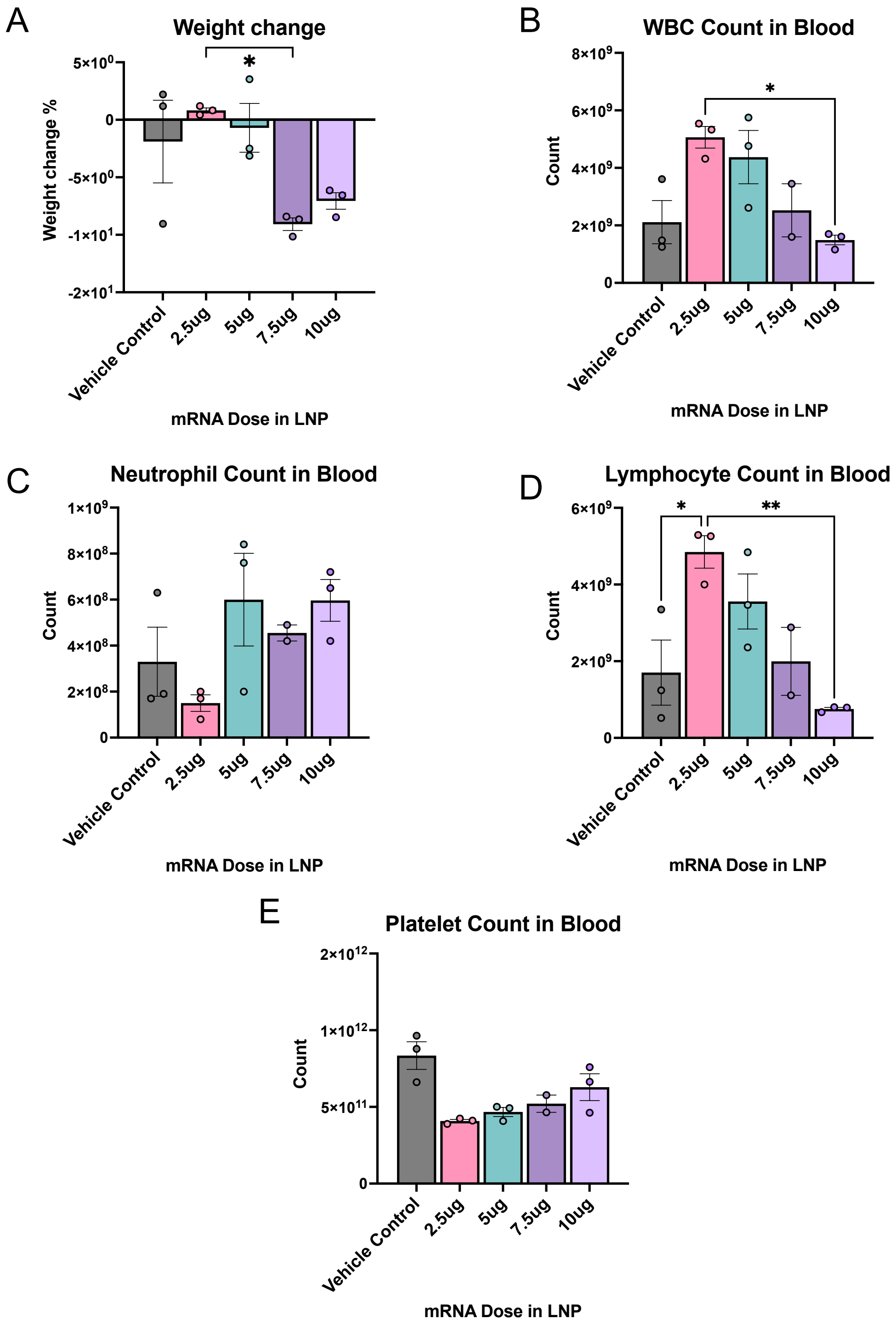


**Supplementary Fig. 1 | Dose-dependent toxicity of intratracheally administered LNPs.** cKK-E12 LNPs were instilled intratracheally into mice at doses of 2.5μg – 10μg of mRNA in LNPs and 24 hours later, the blood and BAL fluid was harvested and analyzed. In mice, cKK-E12 LNPs dose-dependently (A) induce weight loss, (B) decrease white blood cell count, (C) increase neutrophil count, (D) decrease lymphocyte count, and (E) decrease platelet count.

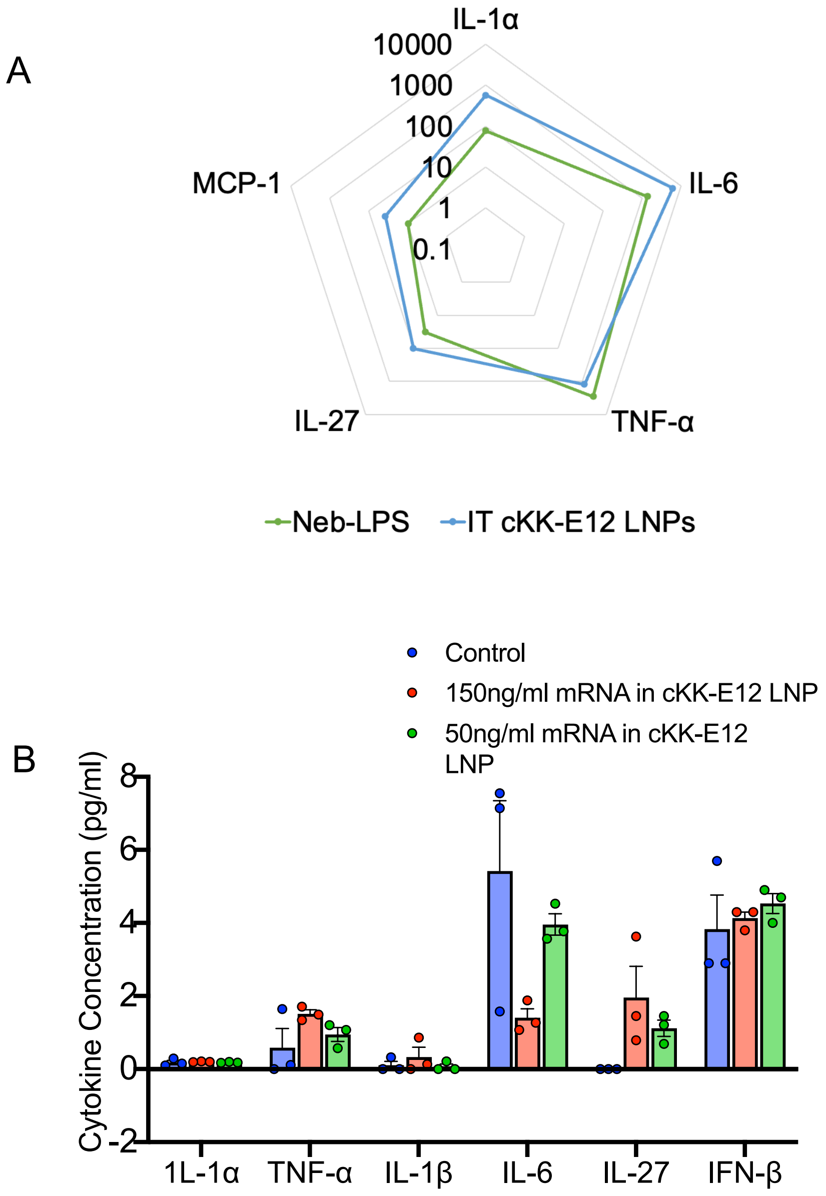


**Supplementary Fig. 2 | Cytokines upregulated by intratracheally administered LNPs and LNPs in MLE-12 cells.** (A) cKK-E12 LNPs were instilled intratracheally into mice at a of 7.5μg of mRNA in LNPs and 2 hours later, the cytokines in the BAL fluid were analyzed and compared to BAL cytokine levels of mice administered with nebulized-LPS 4 hours post-injury (peak of cytokine expression). Mice that had been instilled with LNPs had generally higher cytokine levels in the BAL compared to nebulized-LPS mice. (B) MLE-12 cells were treated with cKK-E12 LNPs and supernatant was analyzed 4 hours later. LNP treatment did not increase cytokine levels above control levels although severe MLE-12 cell death was observed.

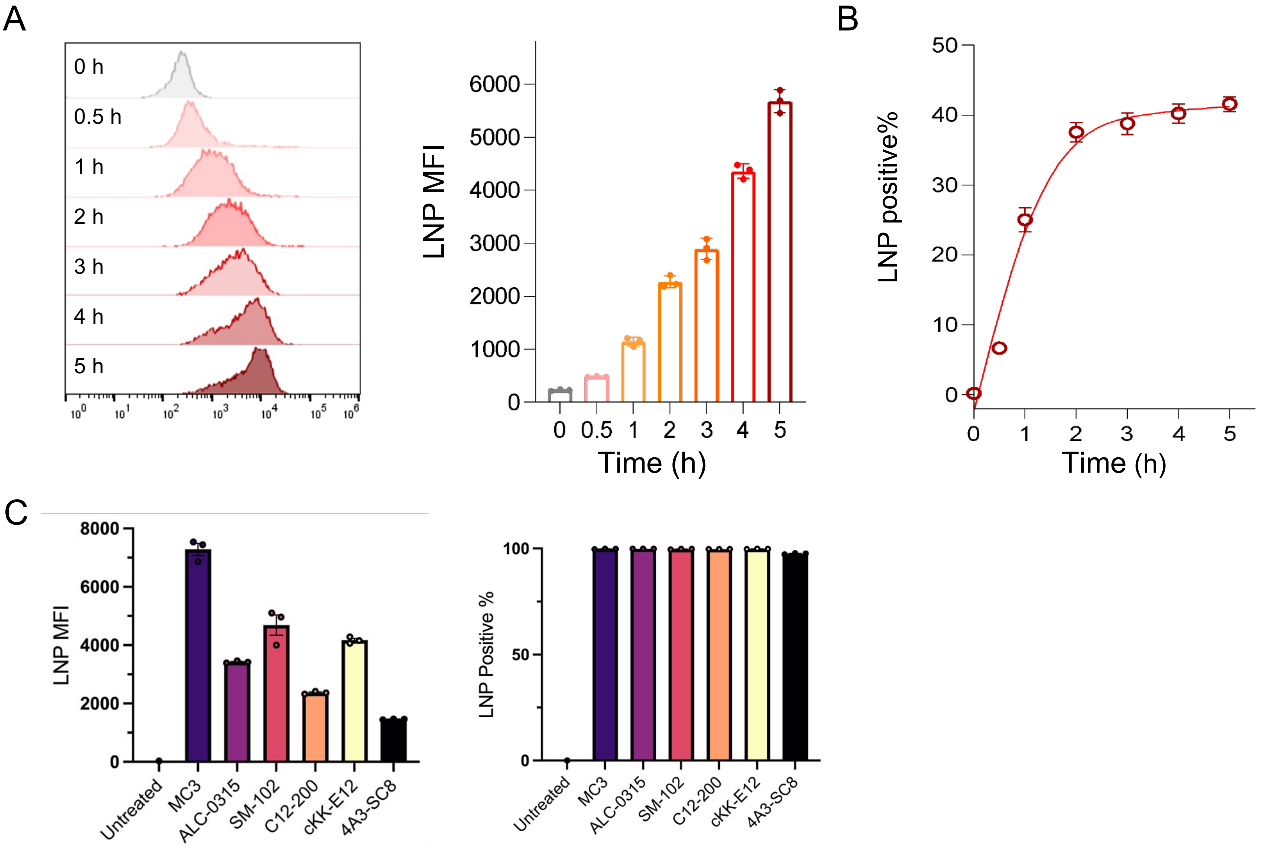


**Supplementary Fig. 3 | LNP uptake in A549$_{GFP1-10}$ and RAW264.7 cells.** To evaluate in vitro LNP uptake, A549$_{GFP1-10}$ and RAW264.7 cells were treated with 18:1 TopFluor PE AF594-labeled LNPs (400 ng/mL). (A) Histogram and mean fluorescence intensity of LNP-treated A549$_{GFP1-10}$ cells. (B) Relative ratio of LNP-internalized cell populations. (C) LNP MFI and uptake % in RAW264.7 cells 6 hours post LNP incubation.

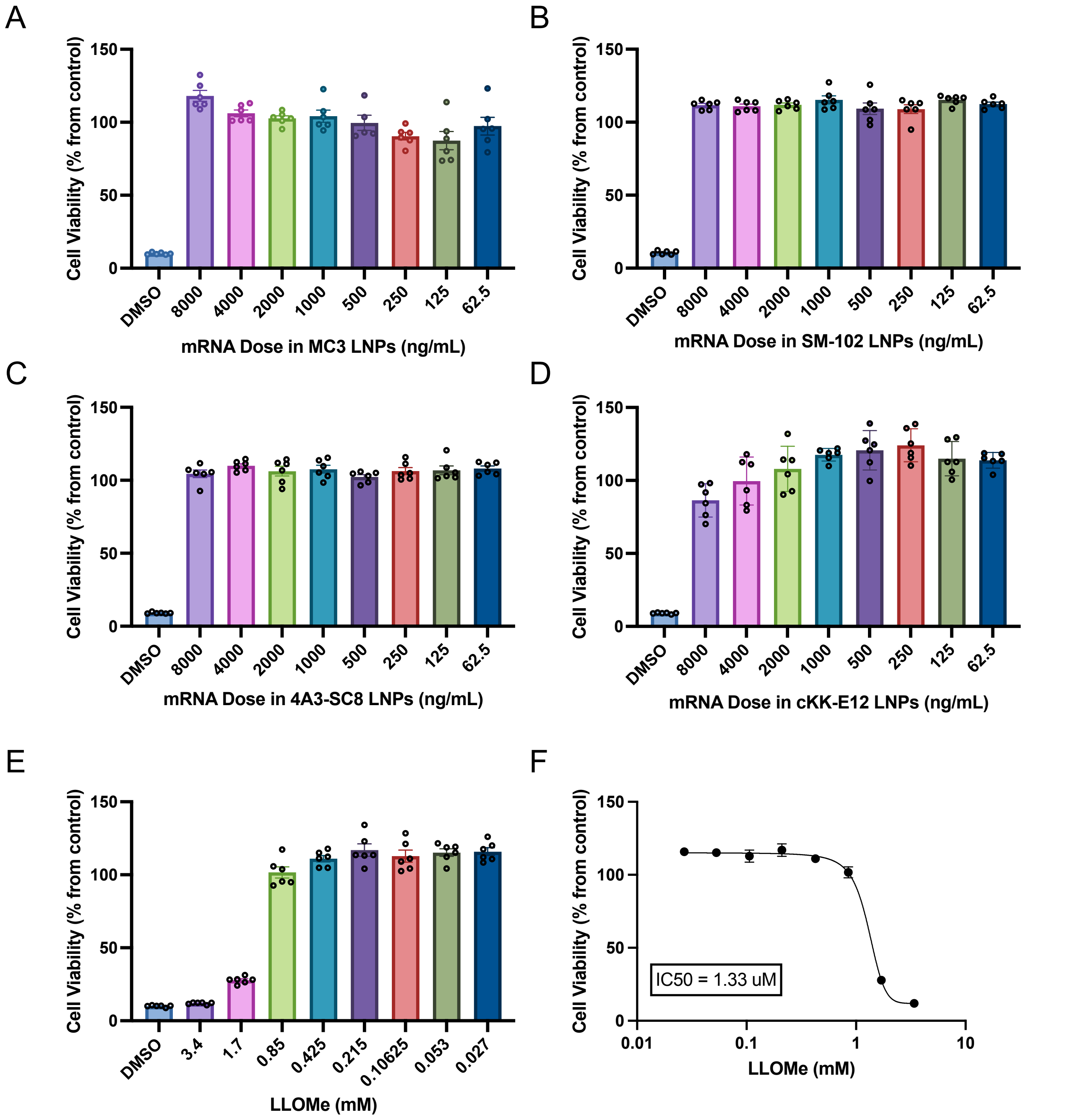


**Supplementary Fig. 4 | Cell viability in RAW264.7 cells.** To evaluate in vitro LNP toxicity, RAW264.7 cells were treated with either MC3, SM-102, 4A3-SC8, or cKK-E12 LNPs (A-D) or LLOMe, a lysosomotropic agent (E, F). After 6 hours, cck8 assay was performed to determine cell viability relative to untreated control. cKK-E12 LNPs led to the most significant levels of cell death amongst the LNP formulations tested.

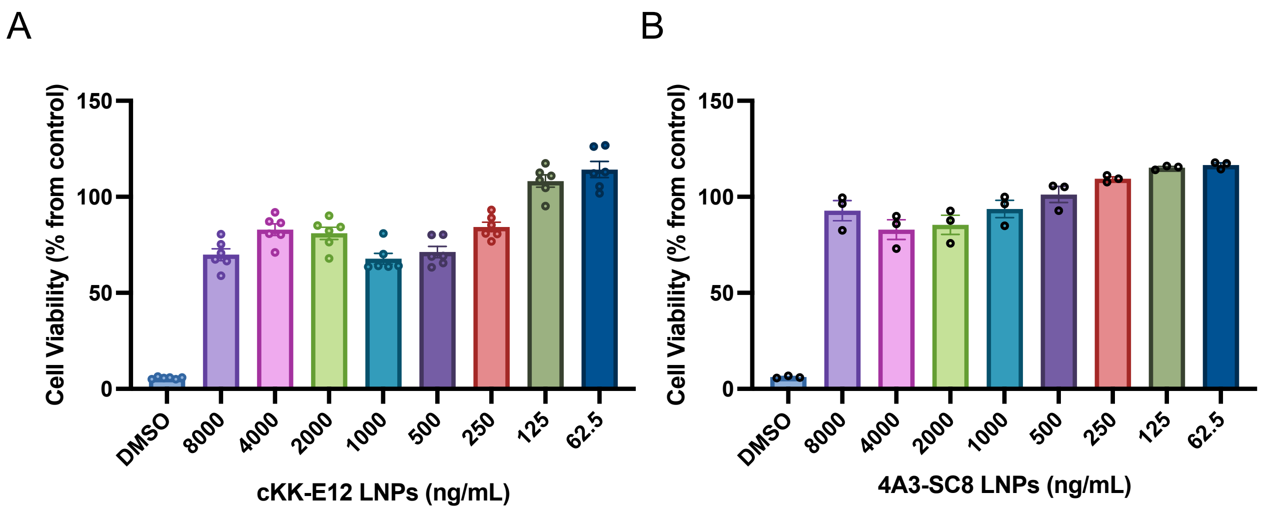


**Supplementary Fig. 5 | Cell viability in A549 cells.** To evaluate in vitro LNP toxicity, A549 cells were treated with either (A) cKK-E12 or (B) 4A3-SC8 LNPs. After 6 hours, cck8 assay was performed to determine cell viability relative to untreated control. cKK-E12 LNPs led to the most significant levels of cell death amongst the LNP formulations tested.

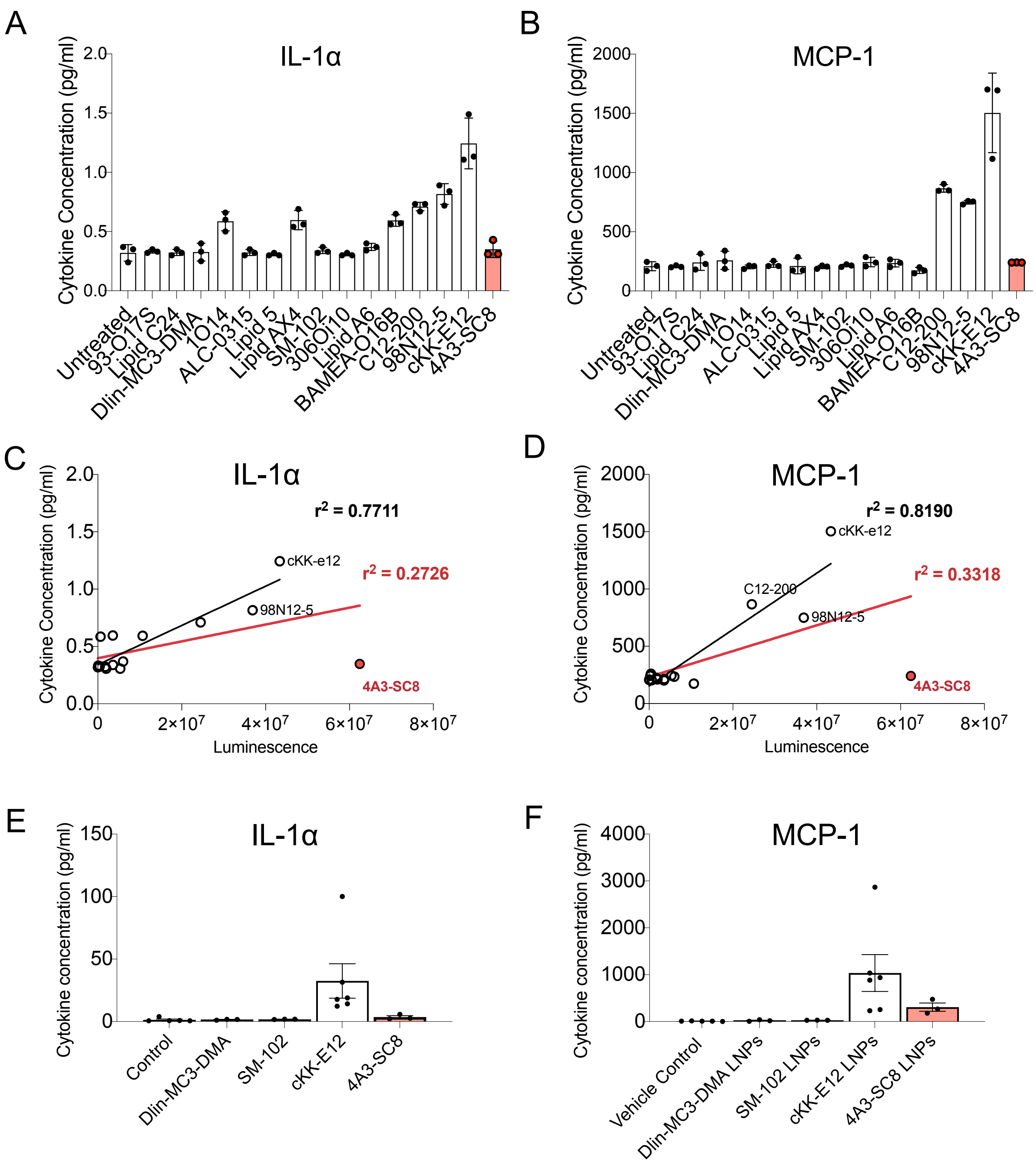


**Supplementary Fig. 6 | LNP-Induced inflammation but also expression of RNA cargo has a positive correlation with ionizable lipid endosomal escape capability.** (A) IL-1α and (B) MCP-1 concentrations 6h after treatment with 15 ionizable lipid LNP formulations in RAW macrophages. *In vitro* (C) IL-1α and (D) MCP-1 concentrations have a positive correlation with luciferase expression. However, 4A3-SC8 LNPs do not increase cytokine levels above control levels. Graphs show the linear regression fits excluding 4A3-SC8 LNPs (black trendlines, (C) $R^2$ = 0.7711 and (D) $R^2$ = 0.8190) versus that including 4A3-SC8 LNPs (red trendlines, (C) $R^2$ = 0.2726 and (I) $R^2$ = 0.3318). After IV LNP injection, the plasma concentrations of (E) IL-1α and (F) MCP-1 follow the trend Dlin-MC3-DMA LNPs< SM-102 LNPs < cKK-E12 LNPs in line with the luciferase expression trend, except 4A3-SC8 LNPs which does not cause significant cytokine upregulation.

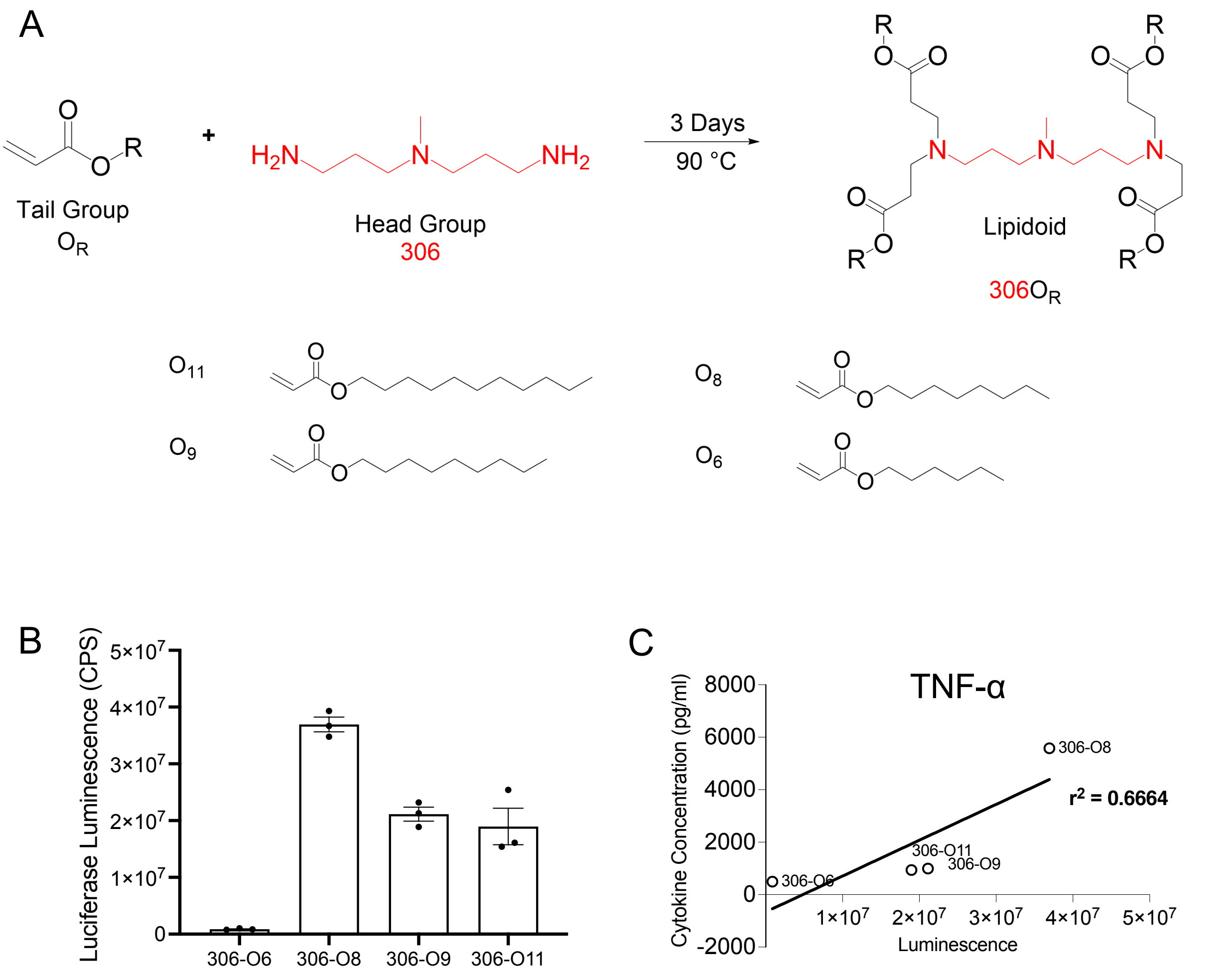


**Supplementary Fig. 7 | LNP-Induced inflammation correlates with RNA expression with a library of lipidoinds with one-carbon differences in tail structures.** (A) 306 series lipidoids with different carbon tail lengths were synthesized as previously described[31] and LNPs were formulated using the same mol fractions in Supplementary Table 1. (B) Luciferase expression of 306-O6, 306-O8, 306-O9 and 306-O11 LNPs in RAW 264.7 macrophages (400ng/ml mRNA in LNPs, 6 hours). (C) Luciferase expression from (B) correlates with TNF-α upregulation.

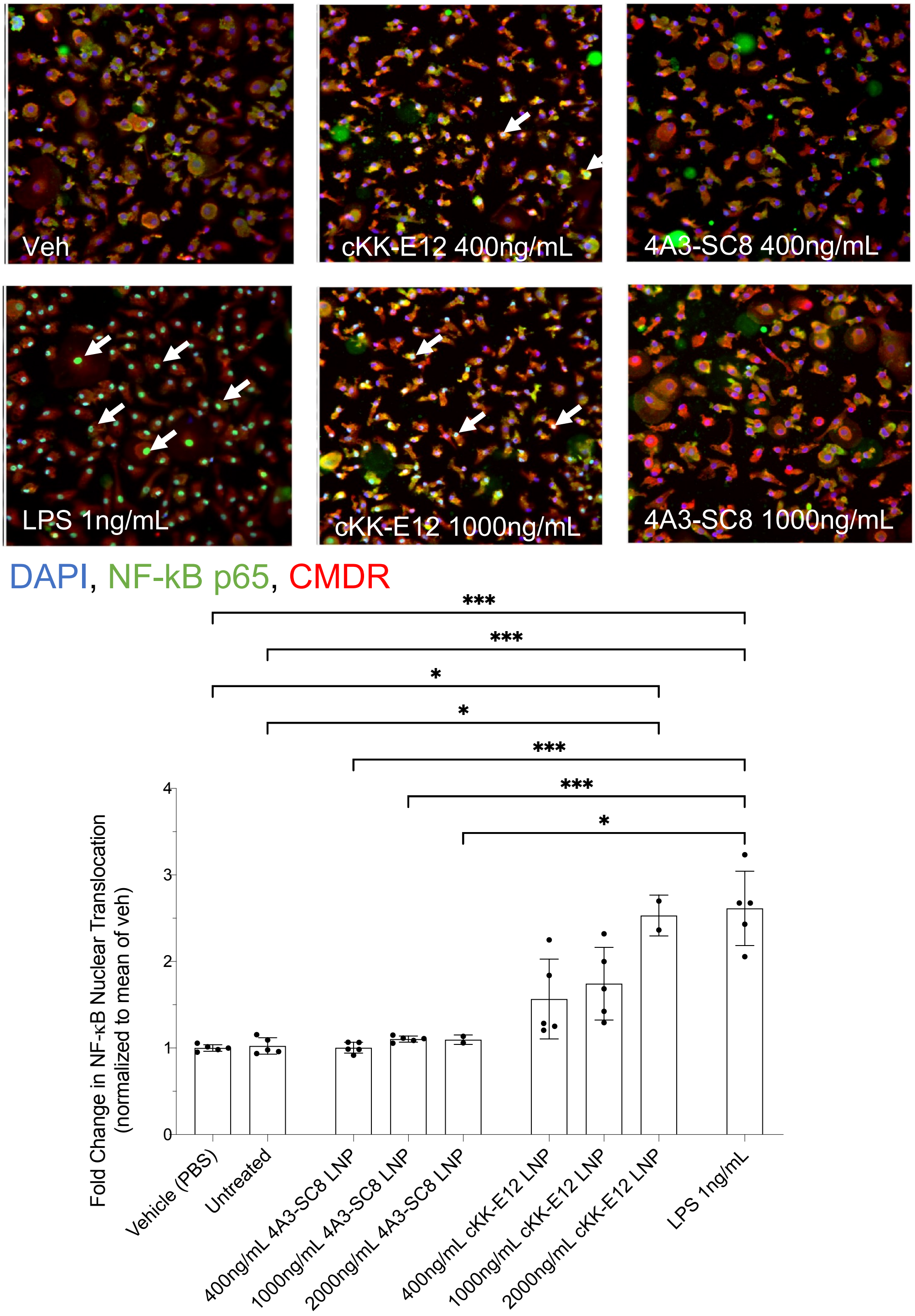


**Supplementary Fig. 8 | Human monocyte-derived macrophage (hMDM) NF-κB response to LNP treatment**. (A) Representative images of NF-κB nuclear translocation in hMDMs treated with vehicle (1 X PBS), cKK-E12 LNPs, 4A3-SC8 LNPs, or LPS (1 ng/mL) for 90 mins. Nuclei are stained with DAPI (blue), cytoplasm with cell mask deep red (CMDR, red). NF-κB p65 (green) is highlighted by white arrows. (B) Fold change in the nuclear translocation ratio (nuclear NF-kB / cytoplasmic NF-kB) in control and treated conditions. hMDM treated with 1000ng/mL cKK-E12 LNPs showed significantly increased NF-kB nuclear translocation while 4A3-SC8 LNPs did not alter NF-kB translocation compared to control levels.

| High Content Imaging Colocalization Analysis Parameters | |
|---|---|
| Condition | Value |
| Ch1: SmoothFactor | 1 |
| Ch1: Thresholding (Fixed) | 500 |
| BackgroundCorrectionCh1 | 50 |
| Object.Ch1.Average Intensity.Ch1 | 0 - 6000 |
| Ch2: SmoothFactor | 3 |
| Ch2: Thresholding (Fixed) | 500 - 850 |
| BackgroundCorrectionCh2 | -255 |
| Ch2: Segmentation (Intensity) | -450 |
| ObjectAreaCh2 | 72.71 - 100000 |
| Object.Average Intensity.Ch2 | 0 - 32767 |
| Ch3: SmoothFactor | 2 |
| Ch3: Thresholding (Fixed) | 400 |
| BackgroundCorrectionCh3 | -255 |
| Ch3: Segmentation (Intensity) | 70 |
| ObjectAreaCh3 | 136.63 - 6065.13 |
| Object.Average Intensity.Ch3 | 0 - 65535 |
| ROI.A.Mask Ch | Channel 1 (DAPI) |
| ROI.A.Target_I | Channel 2 (NF-kB) |
| ROI.B.Mask Ch | Channel 3 (CMDR) |
| ROI.B.Target_I | Channel 2 (NF-kB) |
| ROI.B.Exclude | Channel 1 (DAPI) |

**Supplementary Table 2 |** Parameters used for high content imaging of NF-kB nuclear translocation in primary human MDM.

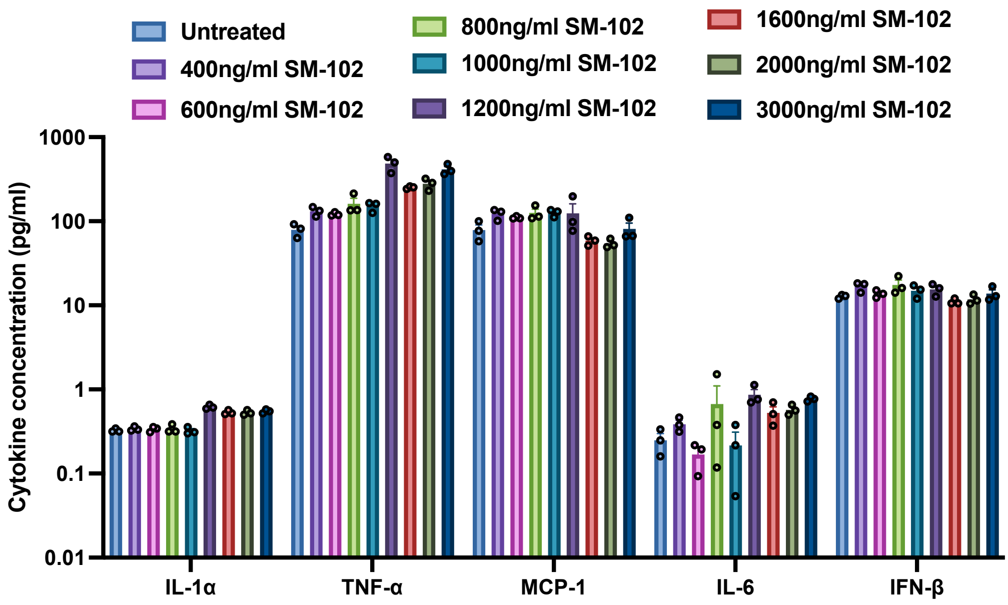


**Supplementary Fig. 9 | SM-102 LNP dose response in RAW264.7 cells.** To evaluate the extent of SM-102 LNP inflammation, RAW264.7 cells were treated with various doses of SM-102 LNPs (400-3000 ng/mL). 6 hours post treatment, supernatant was collected and used for cytokine quantification.

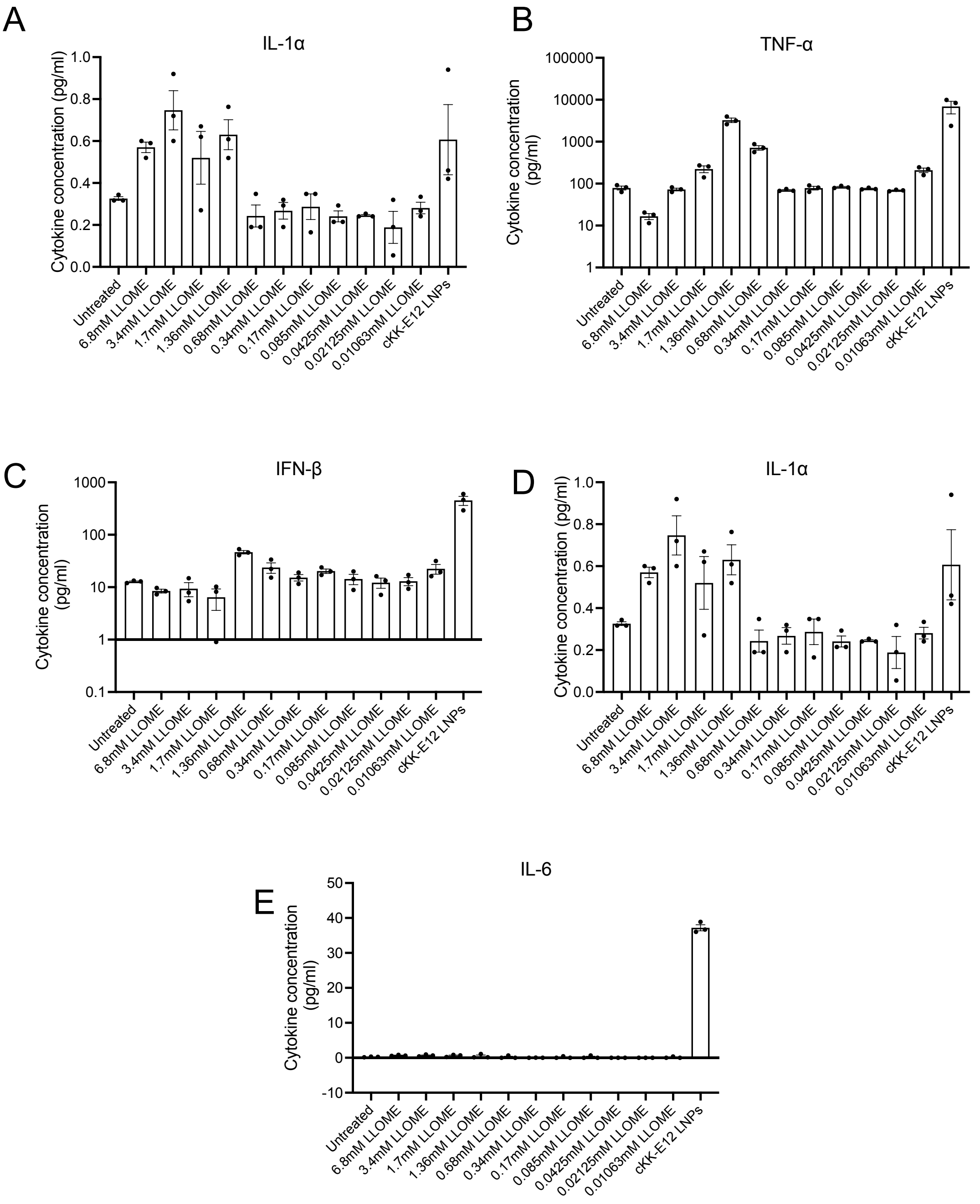


**Supplementary Fig. 10 | LLOME-induced cytokine expression in RAW264.7 cells.** To evaluate the extent of inflammation induced by LLOME compared to cKK-E12 LNPs, RAW264.7 cells were treated with various doses of LLOME (0.01063-6.8mM) or cKK-E12 LNPs (400ng/ml). 6 hours post-treatment, supernatant was collected and used for cytokine quantification. LLOME upregulates mostly the same cytokines as cKK-E12 LNPs

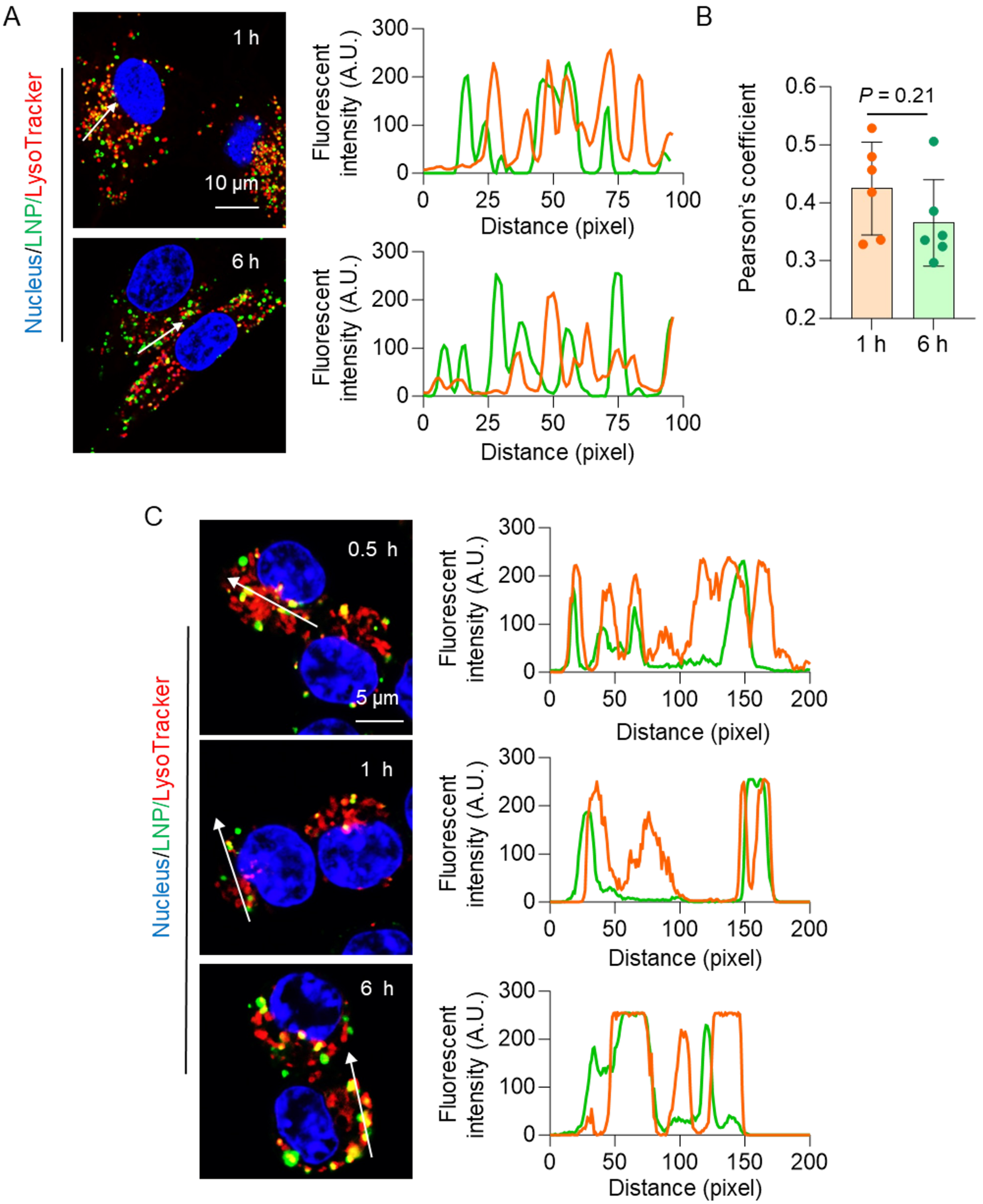


**Supplementary Fig. 11 | Colocalization of LNP and endo-lysosomes.** Cells were treated with 18:1 TopFluor PE AF594-labeled cKK-E12 LNPs (400 ng/mL) for different durations. (A-B) Representative confocal images of endocytosed LNP in A549$_{GFP1\text{-}10}$ cells (A) and Pearson's coefficients (B). (C) Representative confocal images in Raw264.7 cells. Endo-lysosomes were labeled with LysoTracker DeepRed. Nuclei were stained by Hoechst. Line scans show the colocalization of LNPs (green) and endo-lysosomes (red) by pixel. Scale bars: 10 μm (A) and 5 μm (C).

$$\text{Red/Green} = \text{Red MFI/Green MFI}$$

$$\text{Intact endo-lysosome\%} = \frac{\text{Ratio}}{\text{Ratio}_{\text{control}}} \times 100\%$$

$$\text{Ruptured endo-lysosome\%} = 100\% - (\text{Intact endo-lysosome})\%$$

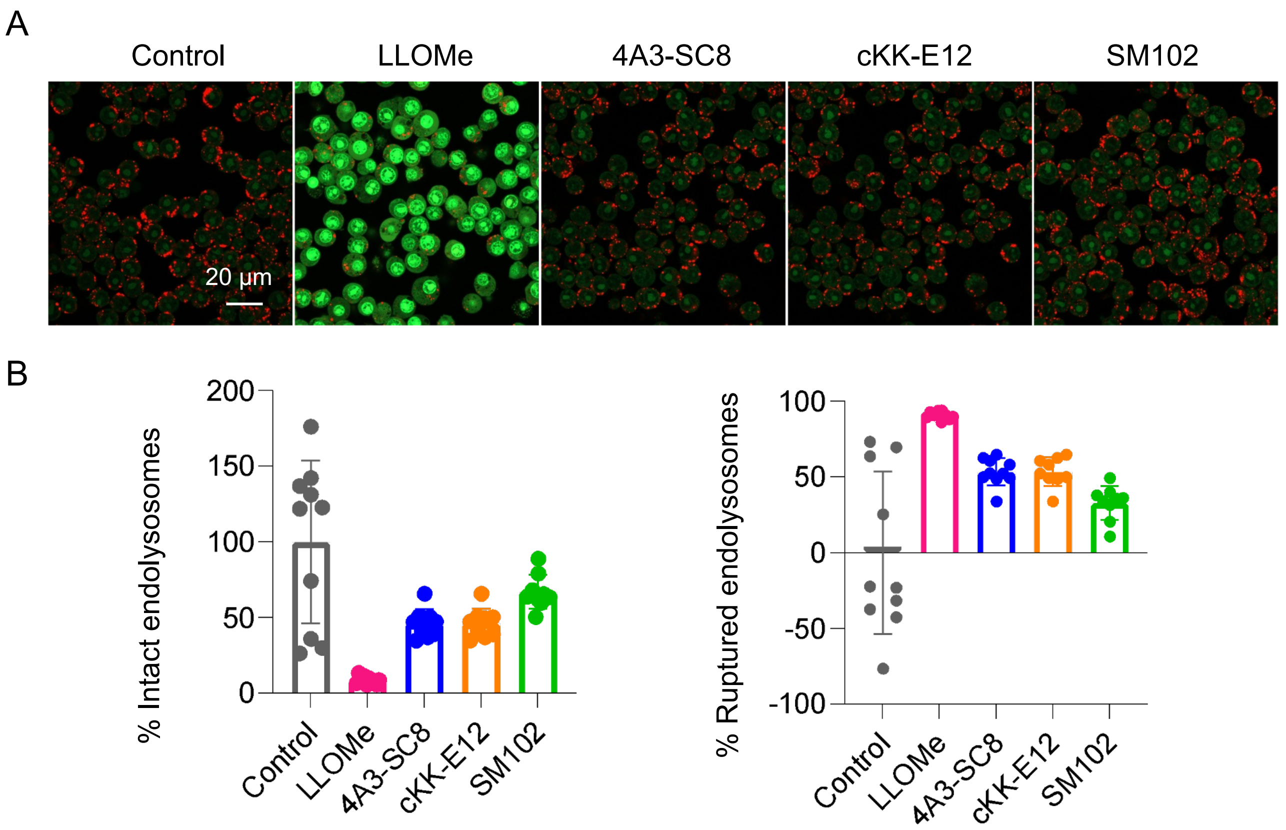


**Supplementary Fig. 12 | Acridine orange imaging in Raw264.7 cells.** (A) Representative confocal images of LNP-treated cells. Scale bar: 20 μm. (B) Quantitative analysis of endo-lysosome damage. Data were normalized to control group. Red = acridine orange in endosomes and green = acridine orange in nucleus/cytosol

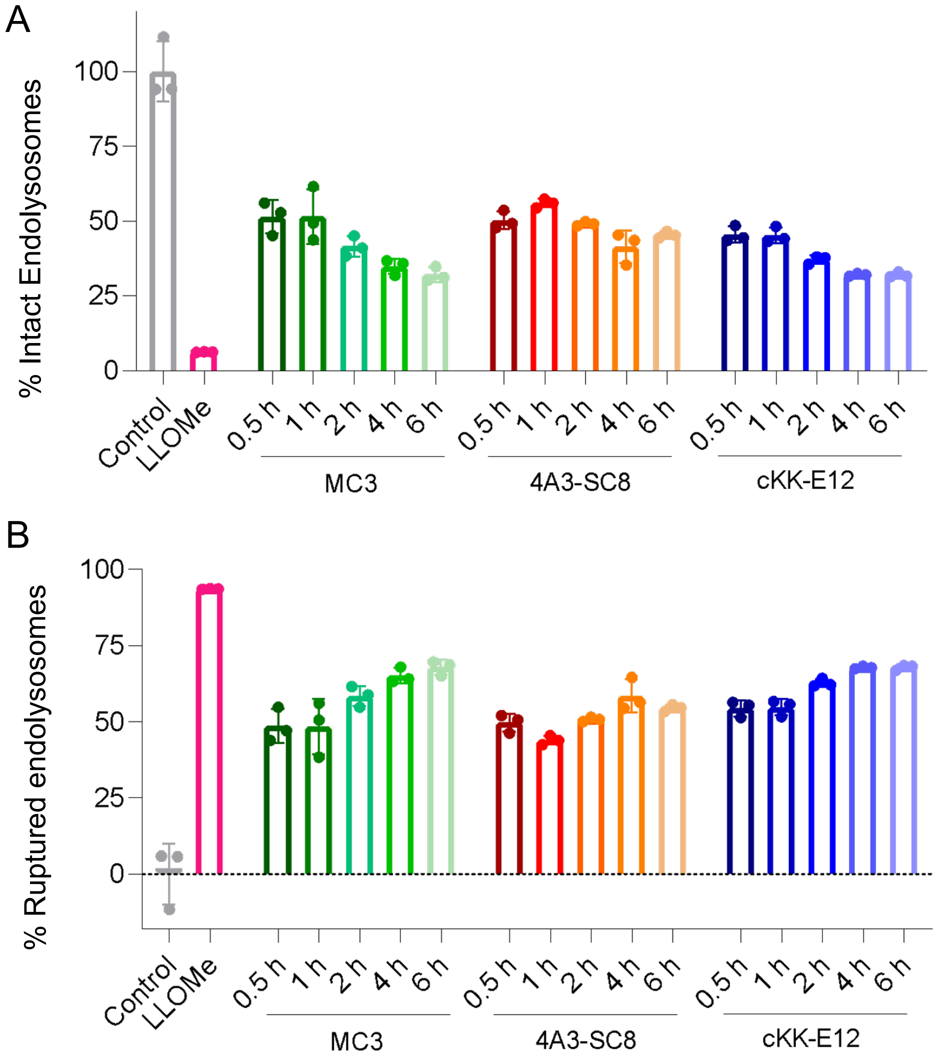


**Supplementary Fig. 13 | Time-dependent endo-lysosome damage in LNP-treated Raw264.7 cells.** (A) Fraction of intact endolysosomes and (B) ruptured endolysosomes after treatment with different LNP formulations at different timepoints. After LNP treatment, cells were stained with acridine orange to test endo-lyososome acidic changes by flow cytometry. Data were normalized to control group.

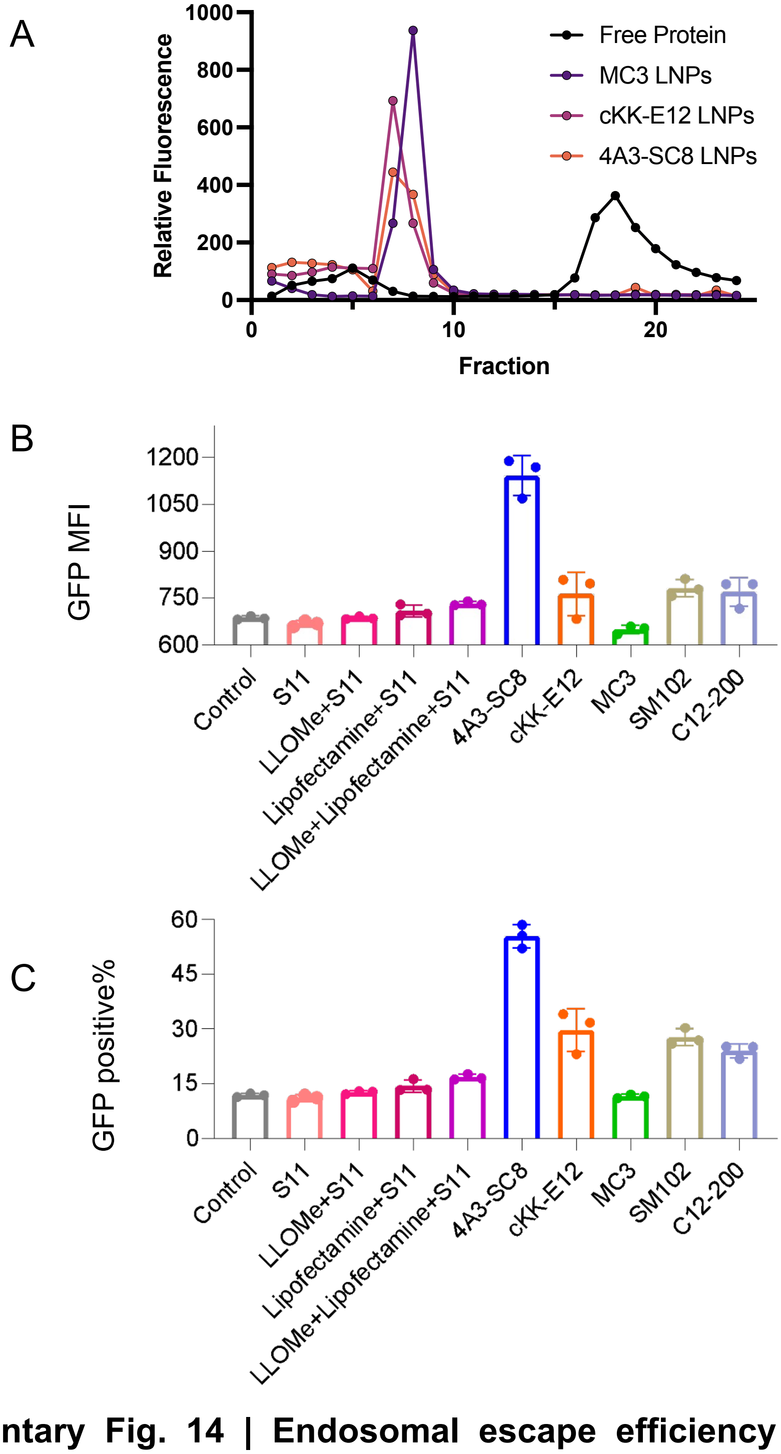


**Supplementary Fig. 14 | Endosomal escape efficiency measured using a split-GFP assay.** A549 GFP$_{1\text{-}10}$ cells were used as split-GFP reporters, which express a nonfluorescent cytosolic fragment of the GFP chromophore (GFP$_{1\text{-}10}$)while the other non-fluorescent peptide fragment S11 was loaded into LNPs. Once LNP-loaded S11 is released from endosomes into the cytosol, the complementation of S11 and GFP$_{1\text{-}10}$ leads to GFP fluorescence. **(A)** Encapsulated split-GFP in LNP was confirmed by running LNPs through a size exclusion chromatograph. A549$_{GFP1\text{-}10}$ cells were treated with free S11 or S11-loaded LNPs for 6 h and then collected for flow cytometry. (B) GFP MFI and (C) fraction of GFP positive cells after treatment with free S11, lipofectamine+S11, LLOME + S11, and LNP-loaded S11. Endosomal escape efficiency generally follows the trend MC3 LNPs < C12-200 LNPs < SM-102 LNPs < cKK-E12 LNPs < 4A3-SC8 LNPs.

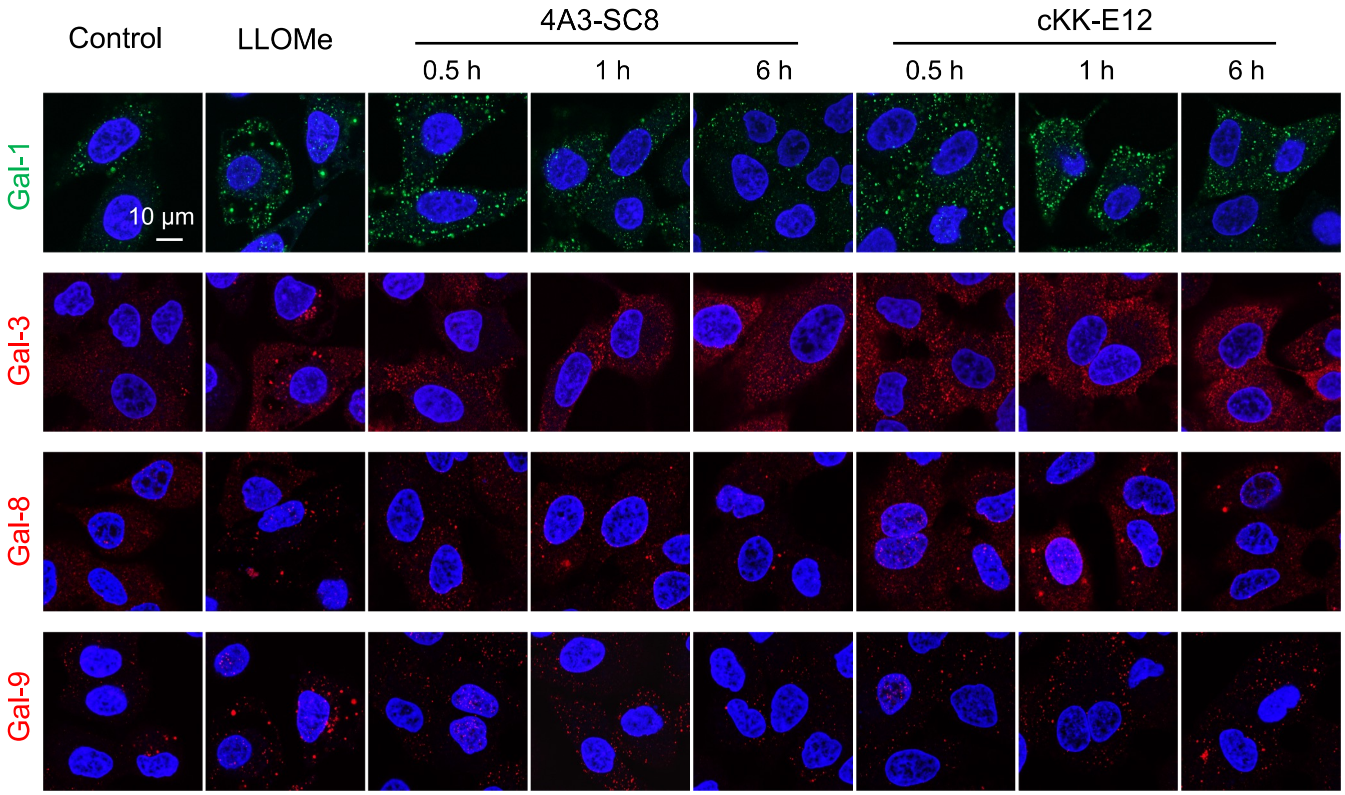


**Supplementary Fig. 15 | Galectin recruitment in A549$_{GFP1\text{-}10}$ cells.** After treatmen with 4A3-SC8 and cKK-E12 LNPs, the intracellular recruitment of gal-1, gal-3, gal-8 and gal-9 were imaged by immunofluorescence. LLOMe was used as a positive control. Scale bar: 10 μm.

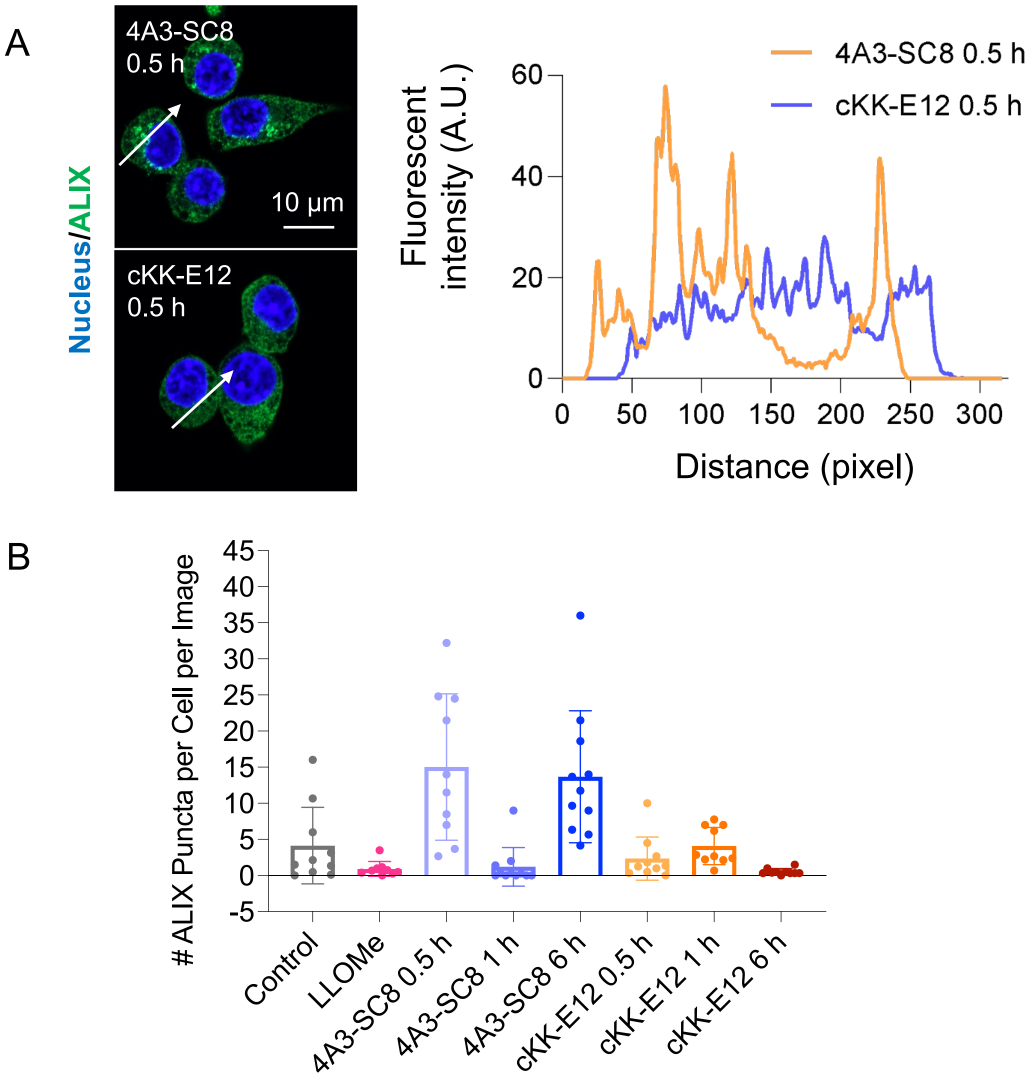


**Supplementary Fig. 16 | Alix puncta in Raw264.7 cells**. (A) Confocal images and line scan of ALIX puncta in LNP-treated cells. Scale bar: 10 μm. 4A3-SC8 LNP-treated cells have significantly more ALIX puncta than cKK-E12 LNP-treated cells. (B) The number of ALIX puncta per cell per image 0.5h, 1h, and 6h post-treatment with 4A3-SC8 or cKK-E12 LNPs.

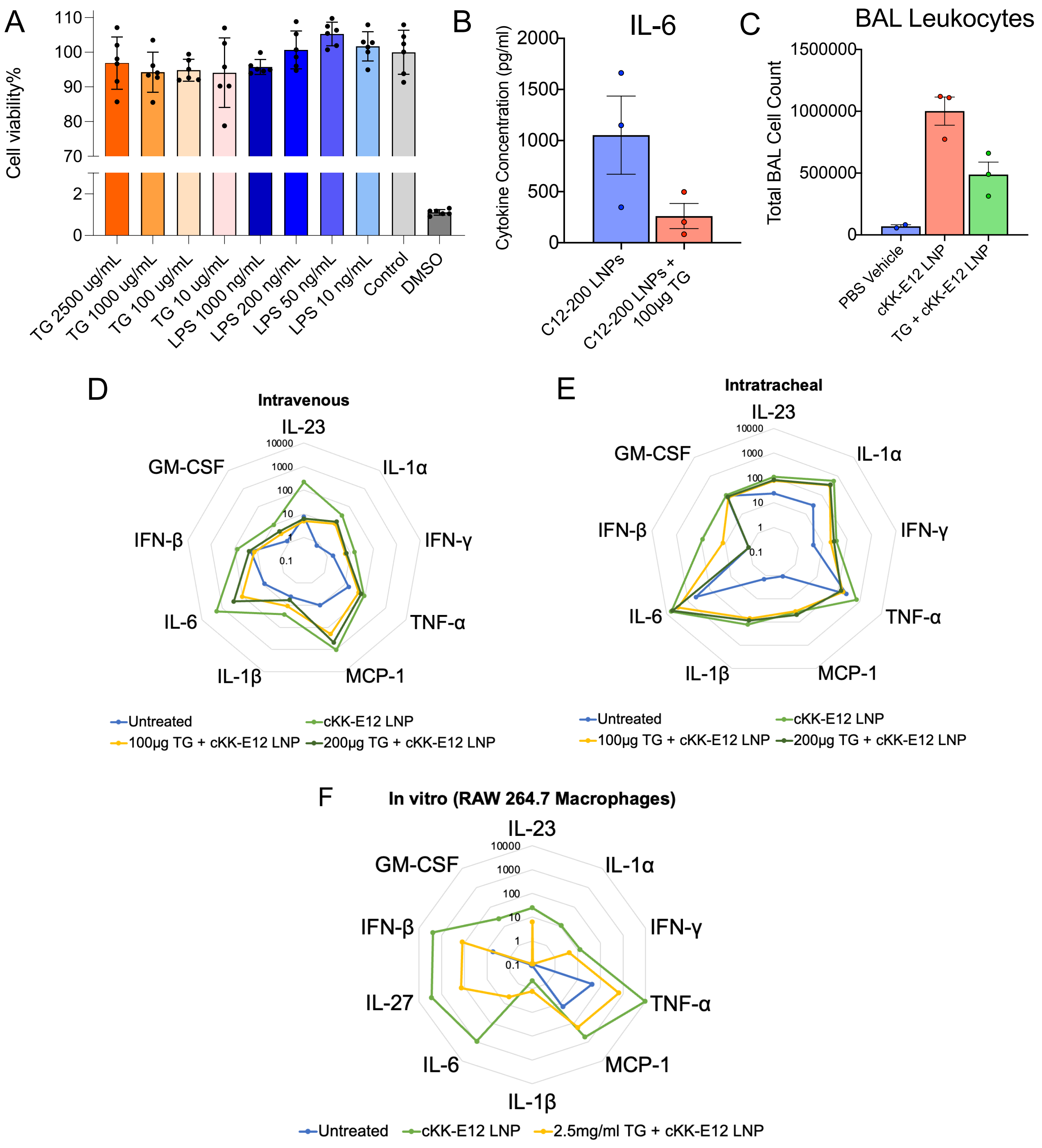


**Supplementary Fig. 17 | TG pre-treatment ameliorates LNP-induced inflammation.** (A) TG treatment does not affect cell viability in MLE-12 cells after 24 hours at doses ranging from 10µg/ml -2500µg/ml. (B) TG pre-treatment reduces the plasma IL-6 levels induced by C12-200 LNPs. TG was intravenously injected 1 hour before C12-200 LNPs and LNPs were allowed to circulate for 2 hours. (C) Intratracheal TG reduces BAL leukocyte levels at longer timepoints. TG was instilled 1 hour before cKK-E12 LNP intratracheal instillation and BAL was harvested 24 hours later LNPs. At different doses, 1 hour TG pre-treatment reduces pro-inflammatory cytokine levels upregulated by LNPs (D) intravenously (7.5µg mRNA in LNP/mouse), (E) intratracheally (7.5µg mRNA in LNP/mouse), and (F) in RAW macrophages (400ng/ml LNP).

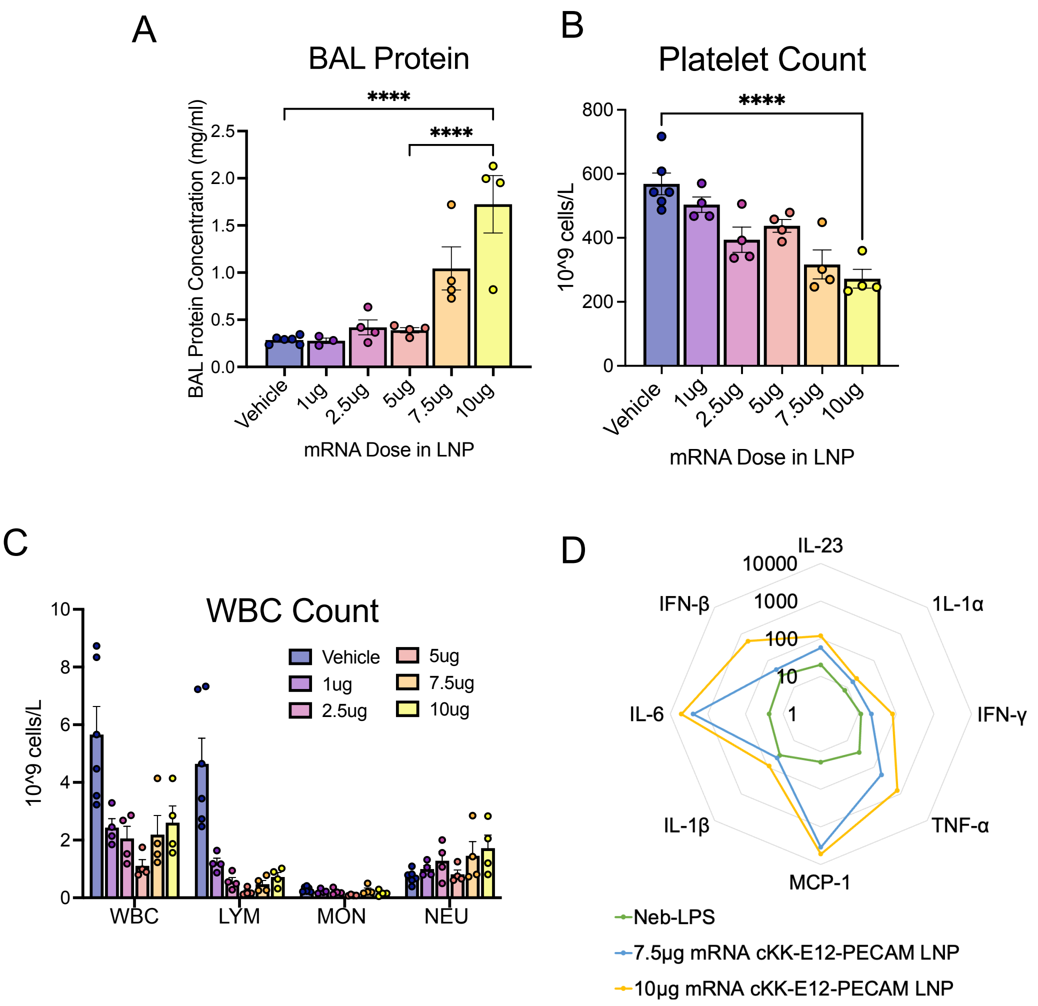


**Supplementary Fig. 18 | cKK-E12 PECAM LNPs exacerbate inflammation in a nebulized-LPS model of ARDS.** In a mouse model of ARDS achieved by administering nebulized-LPS, cKK-E12 LNPs, conjugated to Platelet Endothelial Cell Adhesion Molecule (PECAM) antibodies to target the lung, dose-dependently (A) increase BAL protein levels, (B) decrease blood platelet count, and (C) decrease blood lymphocyte count and increase neutrophil count. (D) cKK-E12 PECAM LNPs increase the plasma levels of pro-inflammatory cytokines by up to three orders of magnitude compared to nebulized-LPS vehicle controls. For A-C, LNPs were injected intravenously 2 hours after nebulized-LPS injury and mice were sacrificed 22 hours after LNP administration. For D, LNPs were injected intravenously 2 hours after nebulized-LPS injury and mice were sacrificed 2 hours after LNP administration. Abbrv: white blood cell (WBC), lymphocyte (LYM), monocyte (MON), and neutrophil (NEU)

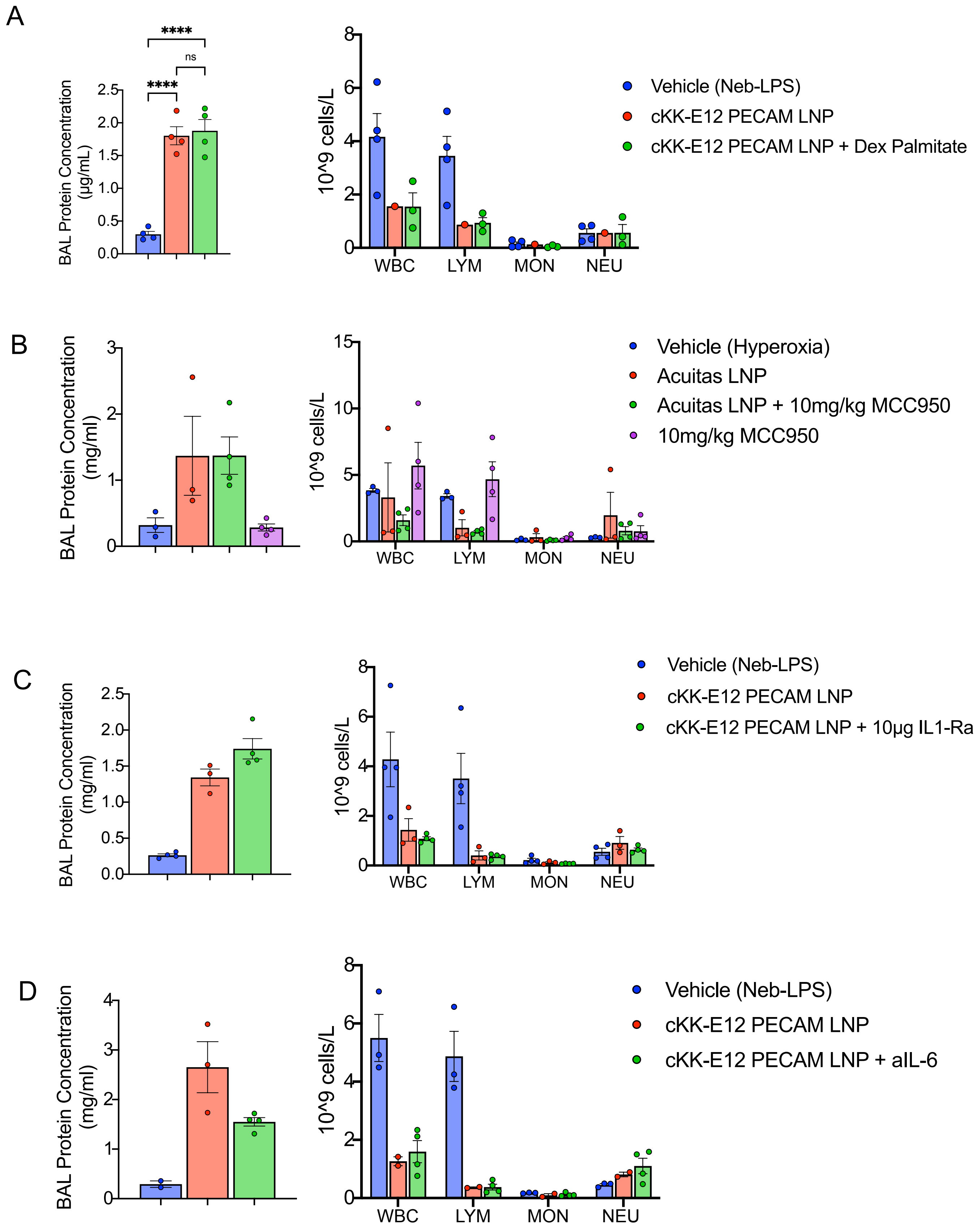
A
****
ns
****
BAL Protein Concentration (µg/mL)
2.5
2.0
1.5
1.0
0.5
0.0
10^9 cells/L
8
6
4
2
0
WBC
LYM
MON
NEU
Vehicle (Neb-LPS)
cKK-E12 PECAM LNP
cKK-E12 PECAM LNP + Dex Palmitate
B
BAL Protein Concentration (mg/ml)
3
2
1
0
10^9 cells/L
15
10
5
0
WBC
LYM
MON
NEU
Vehicle (Hyperoxia)
Acuitas LNP
Acuitas LNP + 10mg/kg MCC950
10mg/kg MCC950
C
BAL Protein Concentration (mg/ml)
2.5
2.0
1.5
1.0
0.5
0.0
10^9 cells/L
8
6
4
2
0
WBC
LYM
MON
NEU
Vehicle (Neb-LPS)
cKK-E12 PECAM LNP
cKK-E12 PECAM LNP + 10µg IL1-Ra
D
BAL Protein Concentration (mg/ml)
4
3
2
1
0
10^9 cells/L
8
6
4
2
0
WBC
LYM
MON
NEU
Vehicle (Neb-LPS)
cKK-E12 PECAM LNP
cKK-E12 PECAM LNP + aIL-6

**Supplementary Fig. 19: Various inhibitors failed to ameliorate exacerbated inflammation induced by LNPs in models of inflammatory lung pathology.** (A) Dexamethasone palmitate loaded into LNPs, (B) 15-minute pre-treatment with MCC950 (10mg/kg), (C) 2-hour pre-treatment with IL-1Ra (10μg / mouse), and (D) 2-hour pre-treatment with IL-6 antibodies (100μg / mouse) did not significantly amerliorate the exacerbated capillary leak into the alveolar space (measured by BAL protein levels), lymphopenia, and neutrophilia induced by LNPs. A, C and D were carried out in the nebulized-LPS model while B was conducted in a hyperoxia model of ARDS achieved by exposing mice to >90% oxygen for 72 hours. For nebulized LPS experiments, LNPs were injected intravenously 2 hours post injury and mice were sacrificed 22 hours after LNP administration. For hyperoxia experiments, LNPs were administered after the 72-hour hyperoxia injury period for a circulation time of 24 hours during which mice were maintained under hyperoxia. Abbrv: white blood cell (WBC), lymphocyte (LYM), monocyte (MON), and neutrophil (NEU).

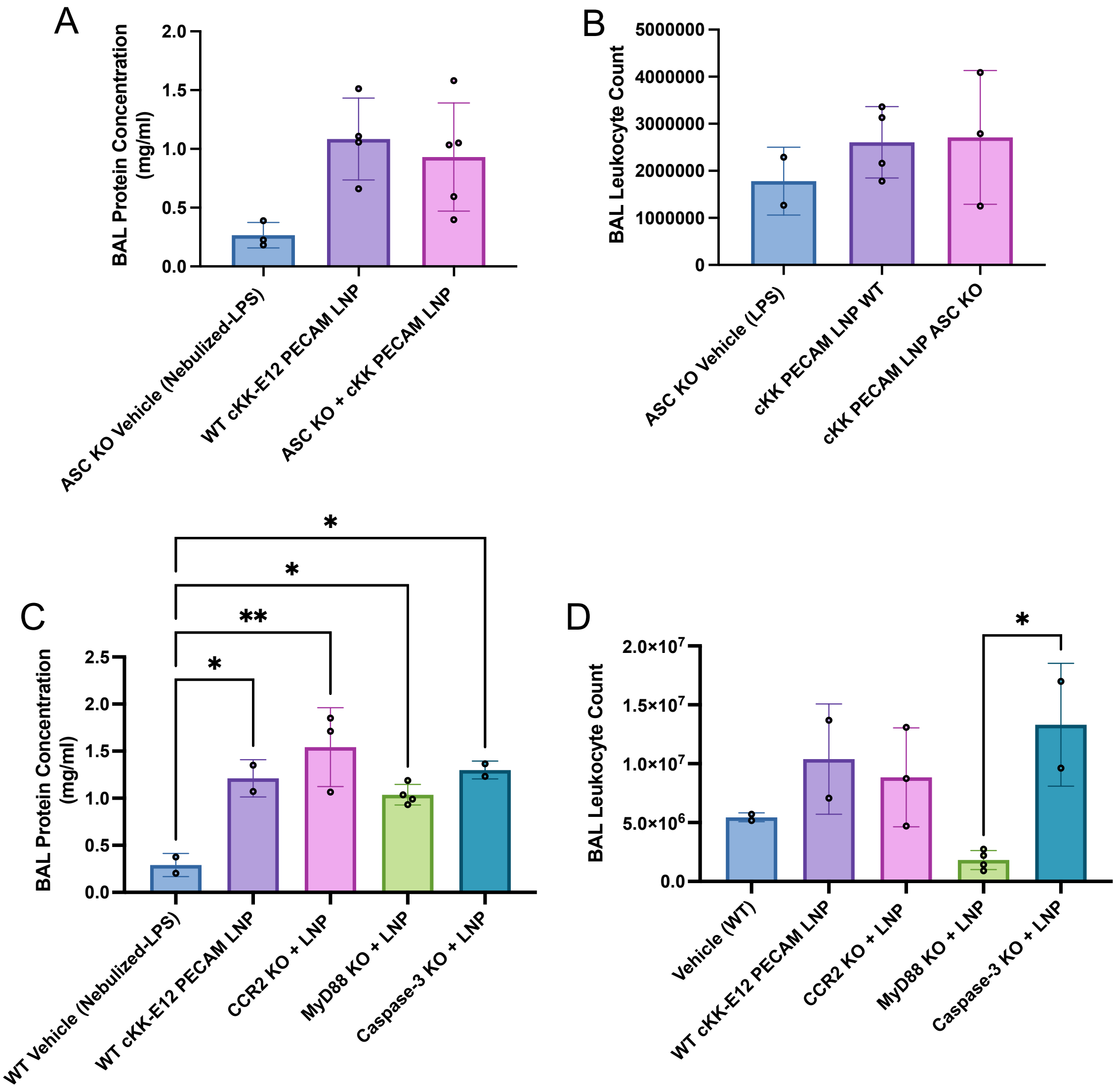


**Supplementary Fig. 20 | LNP-induced inflammation exacerbation persists in various knockout mice.** In LNP-treated ASC knockout mice injured with nebulized-LPS, (A) BAL protein and (B) BAL leukocyte levels are not changed and remain elevated compared to wild - type mice injected with LNPs. (C,D) Nebulized-LPS-injured, CCR2 and caspase-3 knockout mice have similarly elevated BAL protein and leukocyte levels as wildtype mice following LNP injection. MyD88 knockout mice show reduced BAL leukocyte levels compared to wild-type LNP - injected mice and vehicle control mice (nebulized –LPS only). However, MyD88 is vital in the LPS signaling pathway. In all experiments, nebulized-LPS was administered to mice and 4 hours later, PECAM-cKK-E12 LNPs were injected for a circulation period of 20 hours.

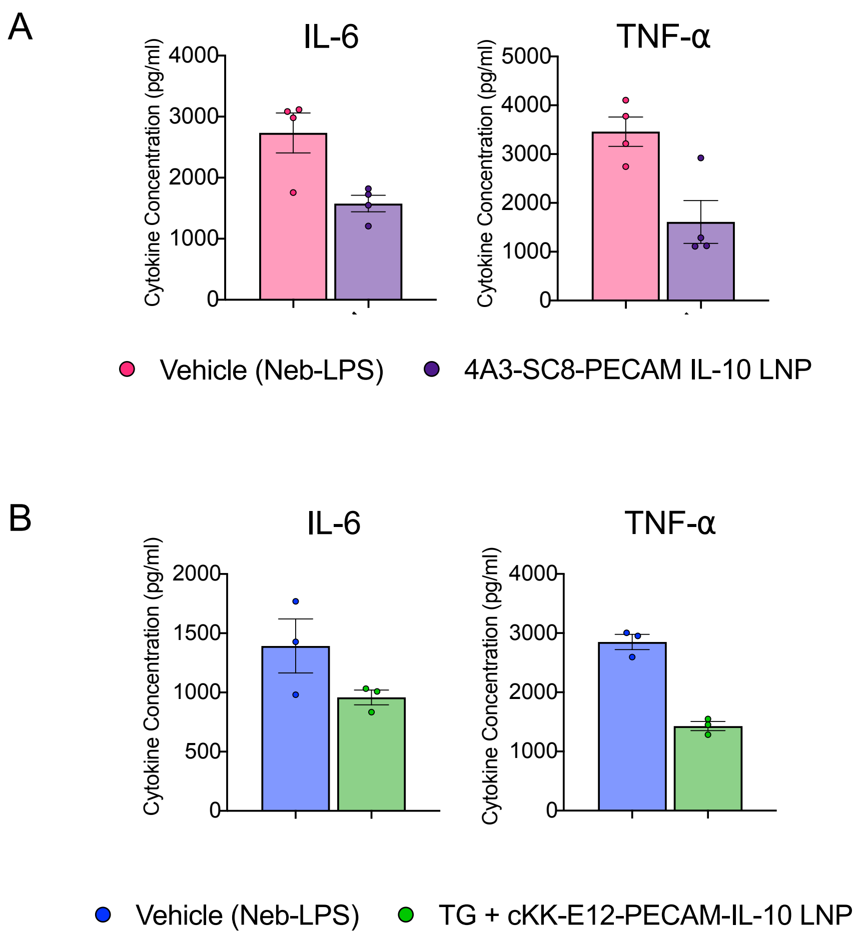


**Supplementary Fig. 21 | IL-10 mRNA-targeted LNPs in Neb-LPS model of ARDS.** To assess the therapeutic efficacy of IL-10 mRNA-LNPs, nebulized LPS was administered to mice along with 0.3mg/kg of mRNA in LNPs. LNPs were surface-modified with PECAM targeting antibody and administered intravenously. (A) IL-10 4A3-SC8 LNPs and (B) TG+ IL-10 cKK-E12 LNPs reduce the BAL concentration of pro-inflammatory cytokines. For cKK-E12 treatment group, 100µg of thiodigalactoside (TG) was administered 1 hour prior to nebulization. 4 hours post nebulized LPS injury and LNP injection, BAL was collected and used for cytokine quantification.

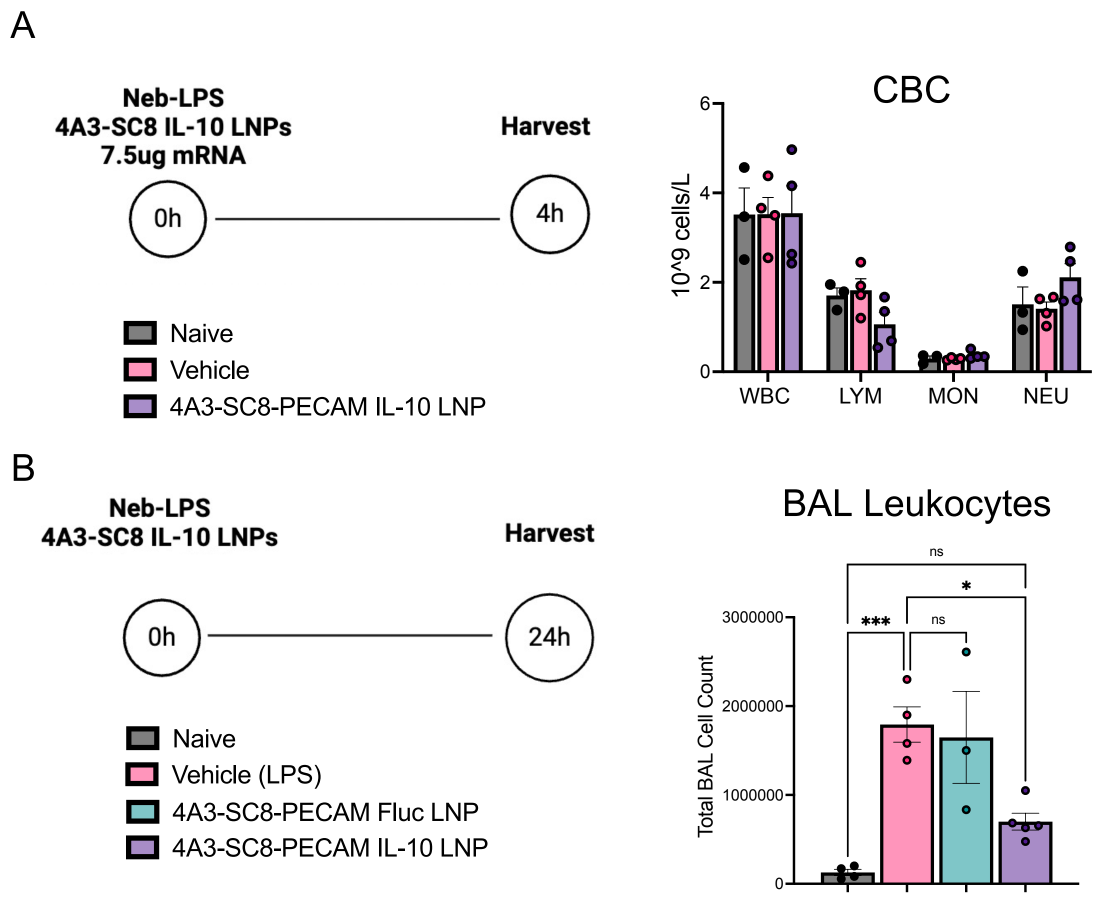


**Supplementary Fig. 22 | 4A3-SC8 PECAM LNPs loaded with IL-10 mRNA ameliorate ARDS phenotypes at longer timepoints.** 4A3-SC8 LNPs loaded with IL-10 (0.3mg/kg mRNA) and conjugated to PECAM (A) do not alter total white blood cell (WBC), lymphocyte (LYM), monocyte (MON), and neutrophil (NEU) counts in the blood compared to control levels and (B) ameliorate leukocyte infiltration into the alveolar space 24 hours after intravenous injection. 4A3-SC8 PECAM LNPs loaded with a model luciferase cargo did not ameliorate ARDS phenotypes but did exacerbate existing inflammation.

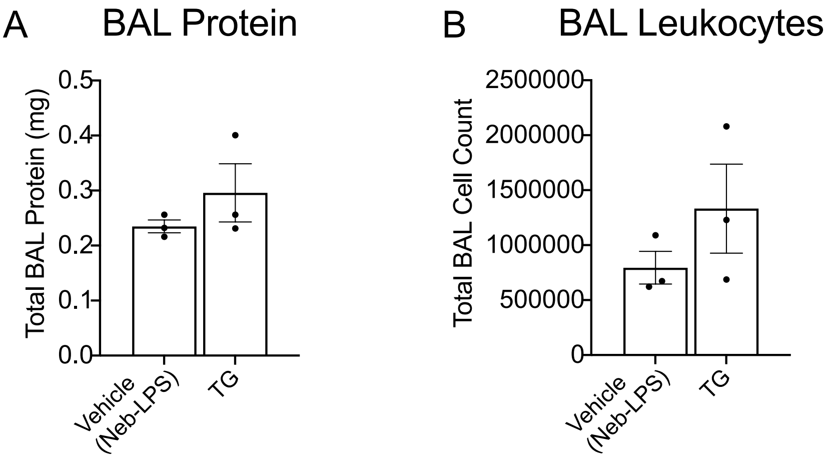


**Supplementary Fig. 23: TG pre-treatment does not affect ARDS phenotypes.** TG administered in the nebulized LPS model does not affect (A) BAL protein levels and (B) BAL leukocyte count. TG was injected intravenously into mice at a dose of 100μg/mouse 1 hour before nebulized LPS injury and mice were sacrificed 4 hours after injury.
